# Evaluation of Commercially Available Glucagon Receptor Antibodies and Glucagon Receptor Expression

**DOI:** 10.1101/2021.12.21.473442

**Authors:** Anna Billeschou Bomholt, Christian Dall Johansen, Sasha A. S. Kjeldsen, Katrine D. Galsgaard, Jens Bager Christensen, Marie Winther-Sørensen, Reza Serizawa, Mads Hornum, Esteban Porrini, Jens Pedersen, Cathrine Ørskov, Lise L. Gluud, Charlotte Mehlin Sørensen, Jens J. Holst, Reidar Albrechtsen, Nicolai J. Wewer Albrechtsen

## Abstract

Glucagon is a key regulator of numerous metabolic functions including glucose, protein and lipid metabolism, and glucagon-based therapies are explored for diabetes, fatty liver disease and obesity. Insight into tissue and cell specific expression of the glucagon receptor (GCGR) is important to understand the biology of glucagon as well as to differentiate between direct and indirect actions of glucagon. However, it has been challenging to accurately localize the GCGR in tissue due to low expression levels and lack of specific methodologies. Immunohistochemistry has frequently been used for GCGR localization, but G-protein-coupled receptors (GPCRs) targeting antibodies are notoriously unreliable. In this study, we systematically evaluated all commercially available GCGR antibodies. Initially, twelve GCGR antibodies were evaluated using HEK293 cells transfected with mouse or human GCGR cDNA. Of the twelve antibodies tested, eleven showed positive staining of GCGR protein from both species. Human liver tissue was investigated using the same GCGR antibodies. Five antibodies failed to stain human liver biopsies (despite explicit claims to the contrary from the vendors). Immunohistochemical (IHC) staining demonstrated positive staining of liver tissue from glucagon receptor knockout (*Gcgr*^−/−^) mice and their wild-type littermates (*Gcgr*^+/+^) with only one out of the twelve available GCGR antibodies. Three antibodies were selected for further evaluation by western blotting and bands corresponding to the predicted size of the GCGR (62 kDa) were identified using two of these. Finally, a single antibody (no. 11) was selected for specific GCGR localization studies in various tissues. In mouse tissue the most intense immunostainings were found in lever, kidney, ileum, heart, and pancreas. Western blotting, performed on liver tissue from *Gcgr^+/+^* and *Gcgr^−/−^* mice, confirmed the specificity of antibody no. 11, detecting a band at high intensity in material from *Gcgr^+/+^*and no bands in liver tissue from *Gcgr^−/−^*mice. Staining of human kidney tissue, with antibody no. 11, showed GCGR localization to the distal tubules. Autoradiography was used as an antibody-independent approach to support the antibody-based findings, revealing specific binding in liver, pancreas, and kidney. As a final approach, RNA-sequencing and single-cell RNA (scRNA)-sequencing were implemented. RNA-sequencing confirmed GCGR presence within liver and kidney tissue. The GCGR was specifically found to be expressed in hepatocytes by scRNA-sequencing and potentially also in collecting and distal tubule cells in the kidney. Our results clearly indicate the liver and the kidneys as the primary targets of glucagon action.

## Introduction

Glucagon is a peptide hormone secreted from pancreatic alpha cells. It is a key regulator of numerous metabolic processes including glucose, protein, and lipid metabolism. These actions are mediated by the GCGR, a GPCR. Knowledge of the GCGR localization and thereby the sites of glucagon action is critical for the understanding of its physiology as well as its potential effects as a therapeutic agent. Besides in the liver, GCGR mRNA has been identified in the adrenal glands^1^, adipose tissue^2^, kidney^1,3^, muscle^4^, and spleen^3^. Northern blot analysis in mice has narrowed the presence of GCGR mRNA down to only the liver and kidney whereas GCGR mRNA by RT-PCR analysis was detected also in the small intestine, lung, brain, and pancreas^5^. In a rat study, GCGR mRNA was detected only in the liver and kidney, but not in adipose tissue, lung, heart, brain, or muscle^6^. These contradictory findings of GCGR localization have left us with the uncertainty as to where the GCGR is expressed. In addition, although mRNA coding for the GCGR may be detected in both rodents^1,2^ and humans^7^, the GCGR mRNA and the corresponding GCGR protein levels at the cell surface do not necessarily correlate since mRNA only represents the transcription and does not guarantee the presence of the mature protein. Therefore, detection of GCGR protein and mapping of the GCGR protein distribution is essential. However, expression levels of GCGR protein in extrahepatic tissues are low and detailed mapping of GCGR protein distribution may therefore require antibody-based approaches including IHC. Antibodies against GPCRs are potentially unreliable as for instance previously reported for the glucagon-like peptide-1 receptor (GLP-1R)^8^. To deal with this problem we systematically evaluated twelve commercially available GCGR antibodies using HEK293 cells transfected with mouse or human GCGR cDNA transcripts and studied liver sections from *Gcgr*^−/−^ and *Gcgr*^+/+^ mice. Based on the evaluation of the twelve available GCGR antibodies, one selected antibody was used to evaluate GCGR localization in mice and humans. Autoradiography, RNA-sequencing, and scRNA-sequencing were used as antibody-independent approaches to support the findings obtained with IHC.

## Material and Methods

### Ethical Approvals

Animal studies were conducted with permission from the Danish Animal Experiments Inspectorate, Ministry of Environment and Food of Denmark, permit 2018-15-0201-01397, and in accordance with the European Union Directive 2010/63/EU and guidelines of Danish legislation governing animal experimentation (1987), and the National Institutes of Health (publication no. 85–23). All studies were approved by the local ethical committee.

Human liver tissue was obtained with approval from the Danish Ethics Committee (H-18052725).

Human kidney tissue was obtained from the European Nephrectomy Biobank (ENBiBA) project (centres from Tenerife and Denmark). Danish Ethics Committee (H-19062684) and Institutional Review Board (IRB) of the Hospital Universitario de Canarias, Spain, approved the project.

### Animals

C57BL/6JRj mice were obtained from Janvier Laboratories (Saint-Berthevin Cedex, France). Glucagon receptor knockout (*Gcgr*^−/−^) mice C57BL/6J^Gcgrtm1Mjch^ were described previously^9^. Homozygotes (*Gcgr*^−/−^) and wild-type (*Gcgr*^+/+^) littermates were used and both were breed with permission from Dr. Maureen Charron and obtained from the local breeding facility. Mice were housed in groups of maximum eight in individually ventilated cages and followed a light cycle of 12 h (lights on 6 AM to 6 PM) with ad libitum access to standard chow (catalogue no. 1319, Altromin Spezialfutter, Lage, Germany) and water. Mice were allowed a minimum of one week of acclimatization before being included in any experiment.

### Antibodies and Reagents

Twelve GCGR antibodies were obtained from a number of different companies (Table 1). The vendors claimed the antibodies to react with GCGR in either human tissue, mouse tissue, or both (Table 1 for specifications). All antibodies were polyclonal and purified from rabbit immune sera. The secondary antibodies used were horseradish peroxidase (HRP)-conjugated, goat anti-rabbit, and rabbit anti-goat immunoglobulins from DAKO A/S (Glostrup, Denmark). Myc-DYKDDDDK Tag monoclonal antibody was obtained from Invitrogen (Naerum, Denmark). Alexa Fluor 546-conjugated goat anti-rabbit IgG (Fab)2 fragment and Alexa Fluor 488 conjugated goat anti-mouse IgG were from Invitrogen (Taastrup, Denmark). All primary antibodies were diluted in PBS with 5% normal goat serum. All other chemicals were from Merck KGaA (Darmstadt, Germany).

**Table 1.** The twelve GCGR antibodies used in the study. All antibodies were polyclonal and purified from rabbit immune sera. Antibody number, name of antibody, vendors, catalogue number, recommended dilution and research resource identifiers (RRID) are listed.

| Antibody number | Name of antibody | Reactivity / target antigen | Vendors, Catalog # | Dilution | RRID |
| --- | --- | --- | --- | --- | --- |
| 1 | GCGR / Glucagon Receptor Antibody LS-C403183 | Human, mouse, rat | LSbio.<br>Cat# LS-C403183 | 1:40 | AB_2890659 |
| 2 | Anti-Glucagon Receptor (extracellular) antibody | Human, mouse, rat | Alomone Labs<br>Cat# AGR-024 | 1:200 | AB_2340975 |
| 3 | Glucagon Receptor antibody | Human, mouse, rat | Biorbyt<br>Cat# orb6093 | 1:400-800 | AB_10920094 |
| 4 | Rabbit Anti-Glucagon Receptor Polyclonal Antibody, Unconjugated | Human, mouse, rat | Bioss<br>Cat# bs-3945R, | 1:200-400 | AB_10856630 |
| 5 | Glucagon Receptor Polyclonal Antibody | Human, mouse, rat | Thermo Fisher Scientific<br>Cat# PA5-50668 | 1:50-200 | AB_2636120 |
| 6 | Rabbit Anti-Human Glucagon Receptor Polyclonal Antibody, Unconjugated | Human | GeneTex<br>Cat# GTX71693 | 8 µg/ml | AB_375890 |
| 7 | Polyclonal Rabbit anti-Human GCGR / Glucagon Receptor Antibody | Human, mouse | LSbio.<br>Cat# LS-C498919 | 1:100 | AB_2890660 |
| 8 | Anti-Mouse Glucagon receptor | Mouse | TriChem (cedarlane)<br>Cat# CL8893AP | 4 µg/ml | AB_2890661 |
| 9 | Glucagon Receptor (extracellular) Polyclonal Antibody | Human, mouse, rat | Thermo Fisher Scientific Cat# PA5-77446 | 1:200 | AB_2735850 |
| 10 | Anti-GCGR polyclonal antibody | Human | Atlas Antibodies<br>Cat# HPA071228 | 1-4 µg/ml | AB_2686368 |
| 11 | Glucagon receptor antibody | Human | Abcam<br>Cat# ab75240 | 5 µg/ml | AB_1523687 |
| 12 | Anti-Glucagon Receptor antibody | Human | Abcam<br>Cat# ab188743 | 13 µg/ml | AB_2890662 |

**Table 2.** An overview of the reactivity of the twelve antibodies, positive or negative glucagon receptor (GCGR) staining on intra- and extracellular domains on transfected HEK293 cells, mouse liver tissue, and human liver tissue. In addition, staining intensity scores between 0 and 3 of liver tissues from both the glucagon receptor wildtype (Gcgr^+/+^) and glucagon receptor knockout (Gcgr^−/−^) mice are presented, the higher the score, the more receptor-antibody binding.

| Antibody number | Reactivity /target antigen, according to vendor: | Intra- and extracellular antibody binding of mouse GCGR transfected HEK293 cells. | Intra- and extracellular antibody binding of human GCGR transfected HEK293 cells. | Extracellular antibody binding of human GCGR transfected HEK293 cells. | Antibody staining of mouse liver tissue and intensity: ($Gcgr^{+/+}/Gcgr^{-/-}$) | Antibody staining of human liver tissue |
| --- | --- | --- | --- | --- | --- | --- |
| 1 | Human, mouse, rat GCGR | Yes | Yes | Yes | No (0/0) | No |
| 2 | Human, mouse, rat GCGR | Yes | Yes | Yes | No (0/0) | No |
| 3 | Human, mouse, rat | Yes | Yes | Yes | No (0/0) | Yes |
| 4 | Human, mouse, rat | Yes | Yes | Yes | No (0/0) | Yes |
| 5 | Human, mouse, rat | Yes | Yes | Yes | No (0/0) | No |
| 6 | Human GCGR | Yes | Yes | Yes | No (0/1) | Yes |
| 7 | Human, mouse, rat GCGR | Yes | Yes | Yes | No (0/3) | Yes |
| 8 | Mouse GCGR | Yes | Yes | No | No (0/0) | No |
| 9 | Human, mouse, rat GCGR | Yes | Yes | Yes | No (0/0) | Yes |
| 10 | Human GCGR | Yes | Yes | Yes | No (0/0) | Yes |
| 11 | Human GCGR | Yes | Yes | Yes | Yes (3/1) | Yes |
| 12 | Human GCGR | Yes | Yes | No | No (0/0) | Yes |

**Table 3.**
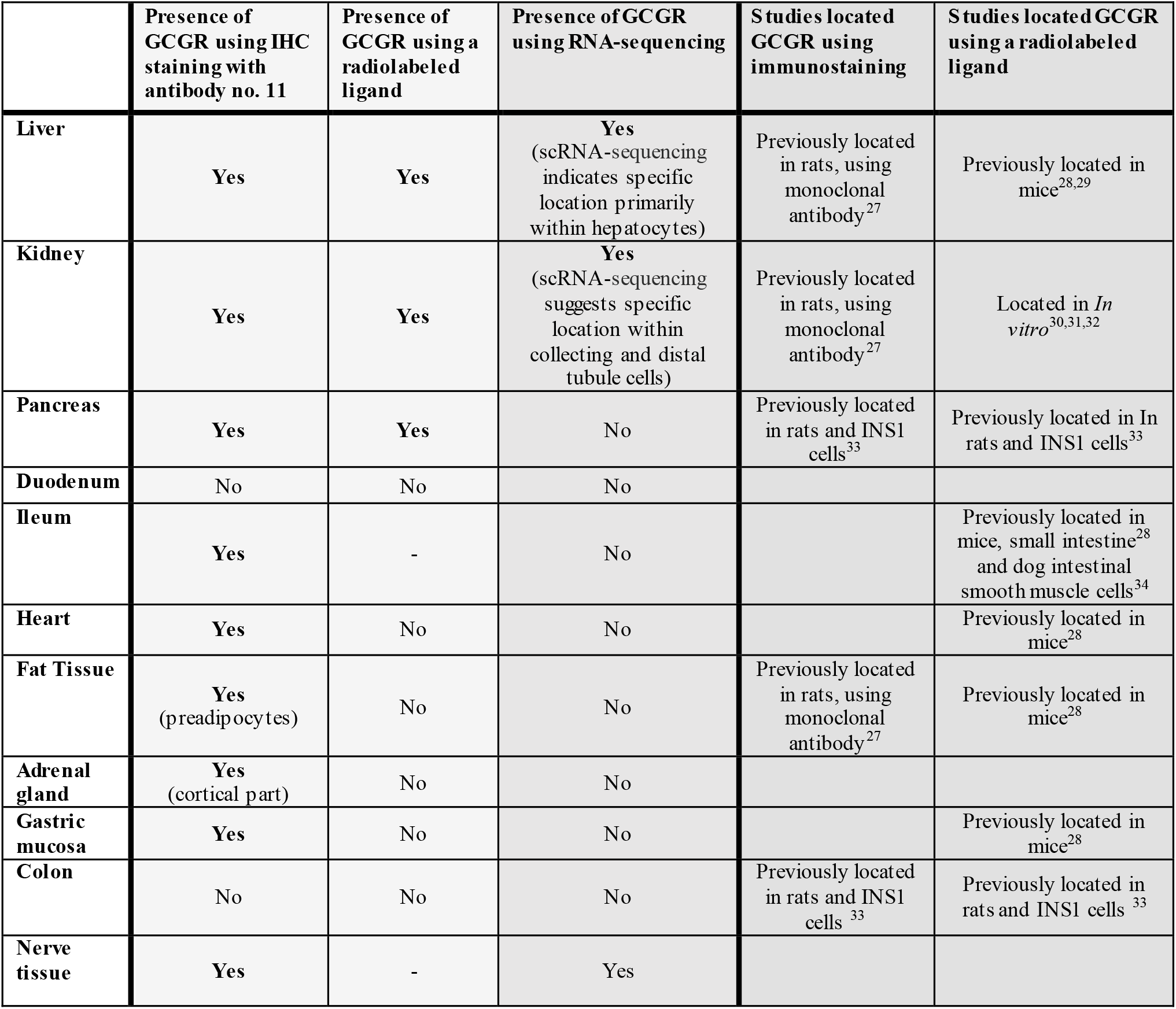
An overview of glucagon receptor (GCGR) expression by antibody and antibody-independent approaches. Demonstrating results from this present study and results previously observed from other studies.

|  | Presence of GCGR using IHC staining with antibody no. 11 | Presence of GCGR using a radiolabeled ligand | Presence of GCGR using RNA-sequencing | Studies located GCGR using immunostaining | Studies located GCGR using a radiolabeled ligand |
| --- | --- | --- | --- | --- | --- |
| **Liver** | Yes | Yes | Yes<br>(scRNA-sequencing indicates specific location primarily within hepatocytes) | Previously located in rats, using monoclonal antibody<sup>27</sup> | Previously located in mice<sup>28,29</sup> |
| **Kidney** | Yes | Yes | Yes<br>(scRNA-sequencing suggests specific location within collecting and distal tubule cells) | Previously located in rats, using monoclonal antibody<sup>27</sup> | Located in *In vitro*<sup>30,31,32</sup> |
| **Pancreas** | Yes | Yes | No | Previously located in rats and INS1 cells<sup>33</sup> | Previously located in In rats and INS1 cells<sup>33</sup> |
| **Duodenum** | No | No | No |  |  |
| **Ileum** | Yes | - | No |  | Previously located in mice, small intestine<sup>28</sup> and dog intestinal smooth muscle cells<sup>34</sup> |
| **Heart** | Yes | No | No |  | Previously located in mice<sup>28</sup> |
| **Fat Tissue** | Yes<br>(preadipocytes) | No | No | Previously located in rats, using monoclonal antibody<sup>27</sup> | Previously located in mice<sup>28</sup> |
| **Adrenal gland** | Yes<br>(cortical part) | No | No |  |  |
| **Gastric mucosa** | Yes | No | No |  | Previously located in mice<sup>28</sup> |
| **Colon** | No | No | No | Previously located in rats and INS1 cells<sup>33</sup> | Previously located in rats and INS1 cells<sup>33</sup> |
| **Nerve tissue** | Yes | - | Yes |  |  |

### Preparation of human and mouse GCGR transfected HEK293 cells

Human embryonic kidney (HEK) 293 cells stably expressing the vitronectin receptor αVβ3 integrin, called 293-VnR, were previously described^10^. Full-length human (RC211179) and mouse GCGR (MR207767) both Myc-DDK-tagged cDNA were obtained from OriGene Technologies, Inc. (Rockville, MD 20850,USA) and inserted into the pcDNA3.1 expression vector. Adherent 293-Vnr cells were transfected with either human or mouse GCGR, using X-tremeGENE9 Transfection Reagent (Roche Applied Science, Hvidovre, Denmark). The cells were cultured in a humidified atmosphere of 95% O_2_ and 5% CO_2_ at 37°C. Blot analysis of total cell lysates was performed to investigate expression levels of the constructs. Mock transfected cells were used as negative controls. Plasmids were sequenced to ensure the right transcript was used (supplementary table 1).

### Immunofluorescence Staining of transfected cells

Visualization of GCGR by immunofluorescence staining was performed as previously described^11^. In brief, the transfected HEK293 cells were cultured for two days, fixated and permeabilized by cold methanol for 10 min at 4°C or for non-permeabilized cells with 4% paraformaldehyde. Fluorescence imaging was performed using an inverted Zeiss Axiovert 220 Apotome system. The images were processed using the Axiovision program (Carl Zeiss, Oberkochen, Germany) and MetaMorph software. For nuclei staining, 4’, 6-diamidino-2-phenylindole (DAPI) was used (Invitrogen; 1:5000).

### Tissue preparation and immunohistochemistry

For preparation of paraffin-embedded tissue samples for chromogenic IHC, tissue from two male and two female *Gcgr*^−/−^ and *Gcgr*^−+/+^ (20-22 weeks of age) mice, liver tissue biopsies from individuals with non-alcoholic steatohepatitis (NASH) and renal tissue biopsies from healthy individuals, were fixed in 10% neutral formalin buffer (BAF-5000-08A, Cell Path Ltd, Powys, United Kingdom) for at least 24 hours at 4°C. Tissue samples from the heart, pancreas, kidney, liver, stomach, duodenum, ileum, colon, epididymal adipose tissue, interscapular brown adipose tissue and adrenal gland were included. Tissues were dehydrated and paraffin-embedded and histological sections of 3 μm were cut. Sections were deparaffinized in xylene and rehydrated using descending alcohol solutions. The sections were transferred in epitope retrieval buffer (Dako, Denmark) and treated in a microwave oven for 2 x 10 min with 20 min pause in between. Endogenous peroxidase was blocked using a hydrogen peroxidase block. Next, a blocking buffer, 5% goat serum in PBS, was added to improve sensitivity by reducing background interference. The sections were incubated with the antibodies overnight at 4°C in a humidified chamber (See Table 1 for application quantity) and biotinylated goat anti-rabbit was added as secondary antibody. To visualize the GCGR positive cells, horseradish peroxidase, alongside its 3,3’diaminobenzidine substrate (Vector Lab, Burlingame, Ca) was added to the sections. For nuclear staining, haematoxylin was used.

### Western blot analysis

Total cellular protein extraction was performed as described previously^12^. Briefly, tissue and or cell line extracted cells were lysed in radioimmunoprecipitation assay buffer containing 50mM Tris/HCl and 140mM NaCl and the pH adjusted to 7.5. The following were added to the buffer:1% Triton X-100, 0.5% SDS,1mM EDTA,1mM PMSF, 1 Tablet of Protease Inhibitor Cocktail (Complete, EDTA-free, Roche). The extract was clarified by centrifugation (14.000 g for 30 min at 4°C). Extracted protein concentration was measured using the Bradford Comassie blue method (Pierce chemical Corp, USA). Protein samples were mixed with reducing sodium dodecyl sulfate (SDS) buffer and separated by 7.5% SDS-polyacrylamide gel electrophoresis, followed by electro-blotting. Non-specific binding was blocked with a 5% nonfat milk solution. The membrane was incubated with primary antibody overnight at 4°C followed by incubation with a horseradish peroxidase-conjugated goat antibody. Protein bands were visualized using enhanced chemiluminescence (Amersham Bio-sciences, Amersham, UK).

### Autoradiography

Non-fasted female C57BL/6JRj mice (13 weeks of age) were anaesthetized by intraperitoneal injection of ketamine/xylazine (0.1 ml/20 g; ketamine 90 mg/kg (Ketaminol Vet.; MSD Animal Health, Madison, NJ, USA); xylazine 10 mg/kg (Rompun Vet.; Bayer Animal Health, Leverkusen, Germany)). When the animals were sufficiently sedated (absence of reflexes), the vena cava was exposed with a midline incision, and the mice received an injection of 3 pmol ^125^I-Glucagon, dissolved in PBS, over a 15 second period (Novo Nordisk, Bagsvaerd, Denmark). Half of the mice also received a 1,000-fold excess (5 nmol) of non-radioactive glucagon in combination with ^125^I-Glucagon (Bachem, Switzerland) in the same injection to test for specific binding. Before injection, 10 μl of the ^125^I-Glucagon stock solution was counted in a γ-counter to determine the amount of radioactivity injected into each animal. After 10 min, the mice were euthanized by cutting the diaphragm to induce pneumothorax and the vascular system was perfused through the left cardiac ventricle (outlet through the right atrium) with saline to ensure removal of blood from the organs. Immediately after, the mice were fixated by perfusion for two minutes with fixative. The small intestine, heart, spleen, liver, kidney and adrenal gland as well as fat and muscle tissue were removed and post fixed in the same solution for 24 h. Tissue samples were embedded in paraffin and sections of 4 μm were coated with 43–45°C Kodak NTB emulsion (VWR, Herlev, Denmark) diluted 1:1 with 43–45°C water, dried, and stored in light-proof boxes at 5°C for 6– 8 weeks. Sections were then developed using Kodak D-19 developer (VWR) for 5 min, dipped 10 times in 0.5% acetic acid, and fixed in 30% sodium thiosulfate for 10 min. Sections were dehydrated, stained with hematoxylin and examined with a light microscope. Images were taken with a camera (Zeiss Axioscope 2 plus, Brock & Michelsen, Birkerød, Denmark) connected to the light microscope.

### Tissue-level gene expression analysis in humans

Gene-level transcript per million (TPM) values were obtained from the GTEx Portal (version 8) and were produced as described on the GTEx documentation page^13^. In brief, RNA-sequencing was done using Illumina TruSeq library construction protocol (non-stranded, polyA+ selection). This consisted of quantification of total RNA using the Quant-iTTM RiboGreen®RNA Assay Kit, polyA selection using oligo dT beads, heat fragmentation, and cDNA synthesis. Following end-repair and poly(A)-tailing, adaptor ligation was performed with Broad Institute-designed indexed adapters. RNA-sequencing libraries were paired-end sequenced (2 x 76 bp) on either an Illumina HiSeq2000 or an Illumina HiSeq2500 instrument according to the manufacturer’s protocols. Reads were aligned to the human genome (GRCh38/hg38) using STAR (version 2.5.3a), based on the GENCODE v26 annotation. TPM values were produced with RNA-SeQC (version 1.1.9) using the *strictMode* flag.

We considered only samples from donors who died fast from natural or violent causes and donors who died unexpectedly with a terminal phase of 1-24 h. The data were filtered for tissues with less than 10 samples and for some tissues with low expression (Cervix Uteri, Bladder, Breast, Ovary, Uterus, Vagina, and Blood), which resulted in 5877 transcriptomic profiles across 22 tissues (Supplementary table 2). Boxplots showing the distribution of TPM values for GCGR across tissues were created using ggplot2^14^ (version 3.3.5) in R (version 4.1.0).

### Cell-level gene expression analysis

The normalized hepatic single-cell RNA-sequencing data were obtained from the GitHub related to MacParland et al., 2018^15^ by use of MacParland’s shiny app. Normalization and clustering were performed by MacParland et al., 2018 using Seurat^16–19^. While MacParland et al., 2018^20^ used t-SNE as dimension reduction for data visualisation, the default Seurat UMAP reduction algorithm was used in this publication. An elbow plot was visually inspected to ensure enough dimensions were included in the UMAP reduction.

The raw renal scRNA-sequencingdata was available from the GitHub related to Liao et al., 2020 publication^21^. The SeuratObject was created differently compared to the Liao et al., 2020 script, as the criteria for *min.cells* and *min.features* were decreased allowing for detection of the lowly expressed GCGR. Quality control, normalization, and clustering were performed as in the original paper. Additionally, the R package Harmony^22,23^ (version 0.1.0) was used to adjust for batch correction and meta-analysis, using settings provided in Liao et al., 2020 “methods” section.

Featureplots, dimplots, and violin plots were performed using built in Seurat functions (FeaturePlot, DimPlot, and VlnPlot, respectively) with parameters available on the GitHub related to this publication. Supplementary table 3 was created using dplyr^24^ (version 1.1.3) in R (version 4.1.0).

## Results

### Evaluation of commercially available GCGR antibodies

#### Antibody staining of transfected HEK293 cells

We initially evaluated whether twelve commercially available antibodies were able to bind to human or mouse GCGR, expressed in HEK293 cells. The two GCGR transcripts harboured both a c-terminal and a cMyc-tag allowing us to double stain cells positive for the GCGR (**Fejl! Henvisningskilde ikke fundet**.). We used permeabilized cells, where the cells have been “pricked” open with methanol, to allow both intracellular as well as extracellular epitope/antibody binding (Figure 1). All GCGR antibodies together with the cMyc-antibody gave positive staining in the permeabilized HEK293 cells transfected with either the mouse or human GCGR transcript. No staining was seen in the controls without primary antibodies (Figure 1). The antibodies were also checked for extracellular staining in HEK293 cells transfected with human GCGR. In this experiment cells were fixed with 4% paraformaldehyde instead of methanol. Ten out of the twelve GCGR antibodies positively stained non-permeabilized HEK293 cells transfected with the human GCGR, while cells stained with antibodies no. 8 and 12 were negative. The GCGR antibodies showed varying efficiency with respect to staining intensity (Figure 2).

**Figure 1.**
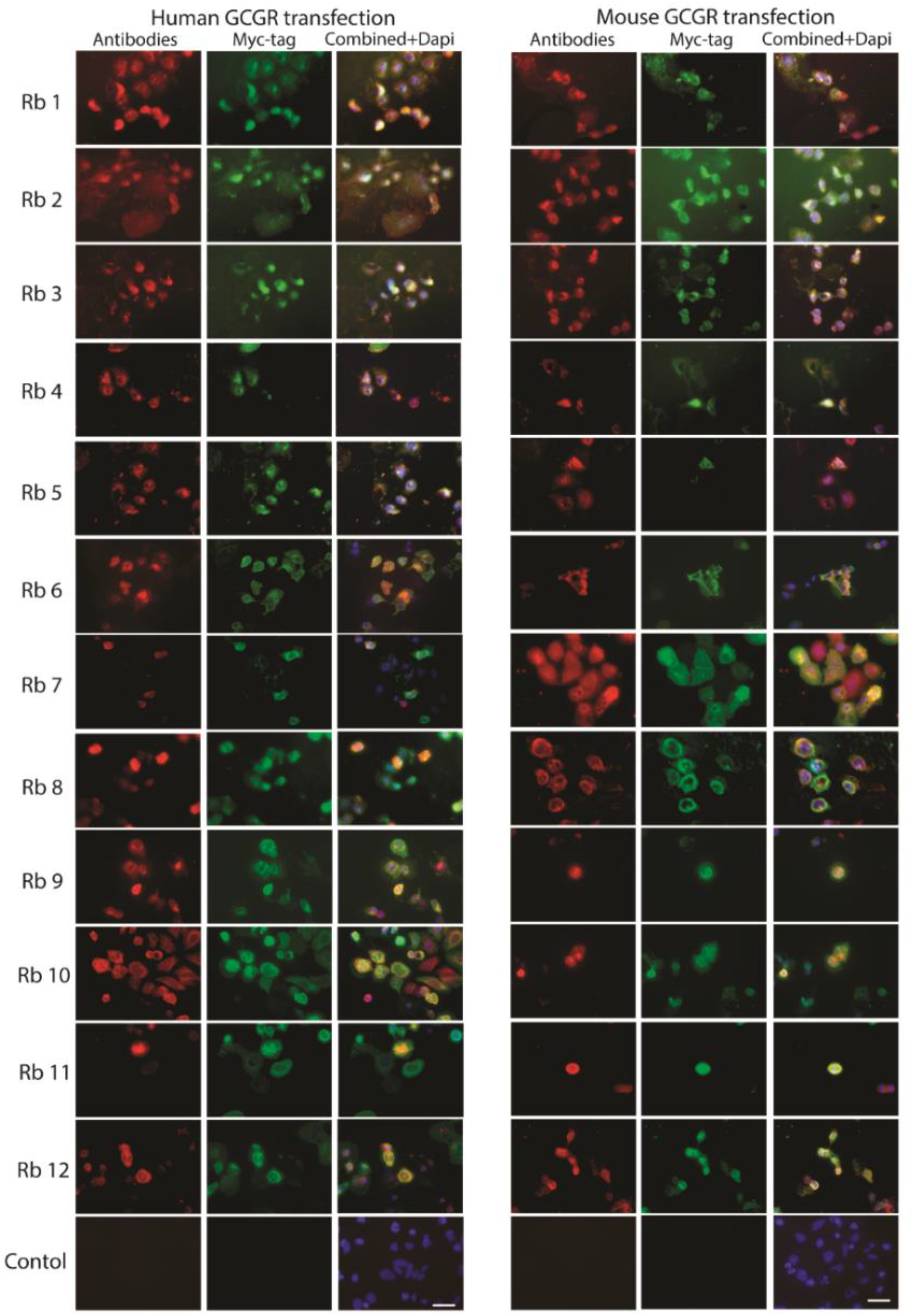
Intra- and extracellular GCGR antibody binding. The twelve antibodies showed varying efficiency in methanol-fixated, permeabilized HEK293 cells transiently transfected with human and mouse GCGR cDNA transcripts. Controls without primary antibodies were used. Red colour indicates GCGR antibody binding, green colour is the Myc-tag on the GCGR and Dapi (blue/purple) indicates nuclei. x85, scale bar = 35 μm. Human GCGR vector: pCMV6-Entry (Cat# PS100001). Mouse GCGR vector: Cat# PS100001.

**Figure 2.**
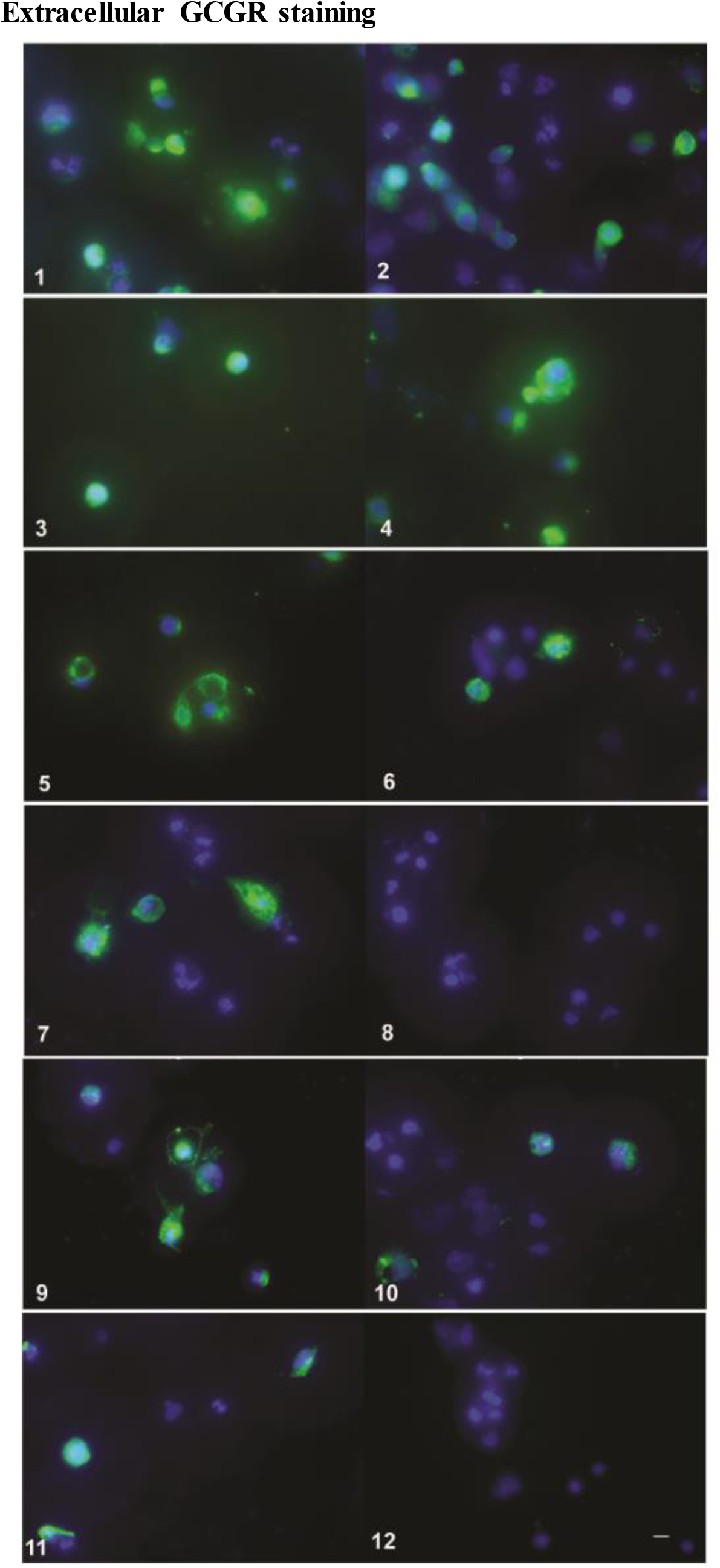
Extracellular GCGR staining. Antibodies no. 8 and 12 did not bind extracellularly. The remaining antibodies showed varying efficiency regarding staining of at the cell membrane surface of non-permeabilized HEK293 cells transiently transfected with human GCGR cDNA transcripts. Blue/purple is dapi (nuclei) and green colour is antibody binding. x150, scale bar = 10 μm. Human GCGR vector: pCMV6-Entry (Cat# PS100001).

#### Antibody staining of liver tissue from Gcgr^+/+^ and Gcgr^−/−^ mice

The twelve commercially available GCGR antibodies (Table 1) were then evaluated using formalin-fixed and paraffin-embedded liver from *Gcgr^+/+^* and *Gcgr^−/−^* mice. Only antibody no. 11 revealed positive IHC staining of liver tissue from the *Gcgr^+/+^* mice. None of the remaining antibodies showed any specific staining of liver tissue from the *Gcgr^+/+^* mice. Antibodies no. 6, 7 and 11 showed unspecific binding in liver tissue from *Gcgr^−/−^* mice with varying intensity (Figure 3).

**Figure 3.**
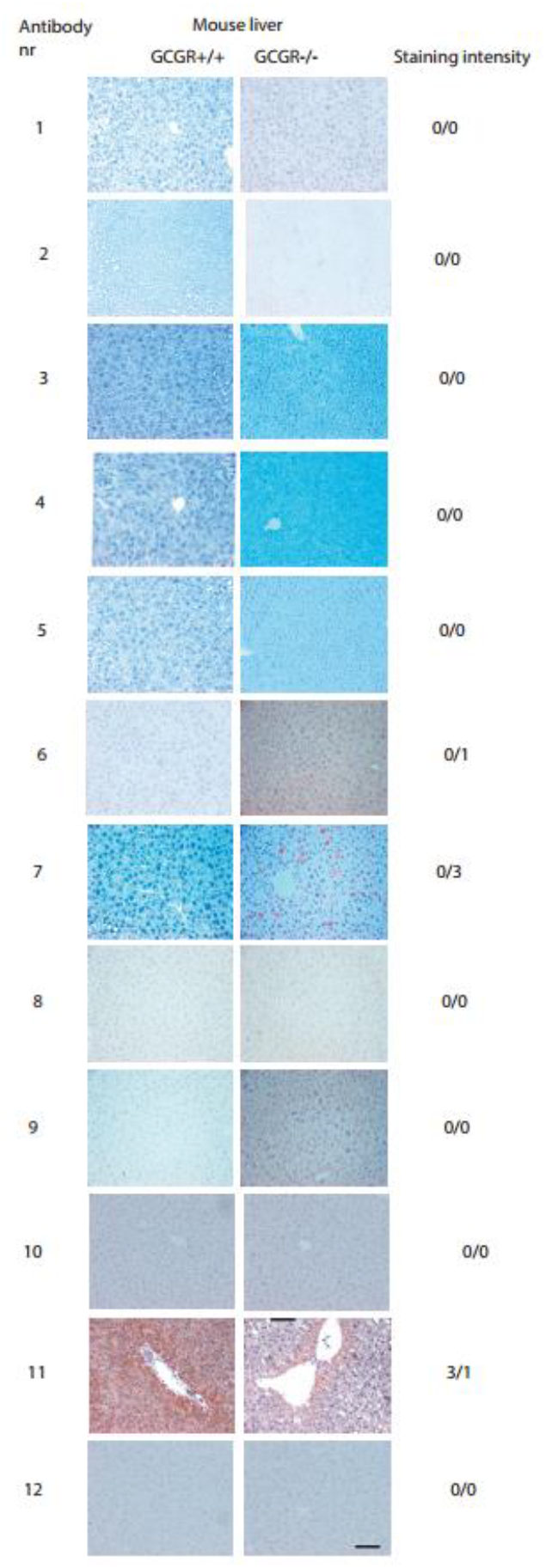
Immunohistochemical staining of liver tissue from either the glucagon receptor wildtype (Gcgr^+/+^) or glucagon receptor knockout (Gcgr^−/−^) female mice, 8 weeks of age, using the 12 GCGR antibodies. Staining intensity scores between 0 and 3 of liver tissues from both the Gcgr^+/+^ and Gcgr^−/−^ mice are presented (^+/+^/^−/−^), the higher the score, the more receptor-antibody binding. X40, scale bar = 50 μm.

#### Antibody staining of human liver tissue

To ensure the applicability of these antibodies for human use, we also evaluated the twelve commercially available GCGR antibodies on paraffin-embedded liver tissue biopsies from patients with NASH. Antibodies no. 1, 2, 5, 8 and 9 did not bind to the human liver tissue, whereas antibodies no. 3, 4, 6, 7, 10, 11 and 12 showed varying staining intensity (Supplementary figure 1).

Based on these current findings we selected three GCGR antibodies (antibodies no. 4, 10 and 11) for further evaluation by western blotting.

#### Comparison of the reactivity of the twelve antibodies

#### Further evaluation of selected antibodies using Western Blotting

To further evaluate the specificity of the antibodies we performed western blotting of extracts of HEK293 cells transfected with either human or mouse GCGR cDNA.

Bands similar to that of the predicted size of the human and mouse GCGR (62kDa) were observed using antibodies no. 10 and 11. Larger bands were detected at high intensity using antibody no. 10 whereas antibody no. 11 appeared to be more specific. Antibody no. 4 detected multiple bands and was not further evaluated (Figure 4). Antibody no. 11 was selected as the most specific. Uncropped western blots are shown in supplementary figure 2.

**Figure 4.**
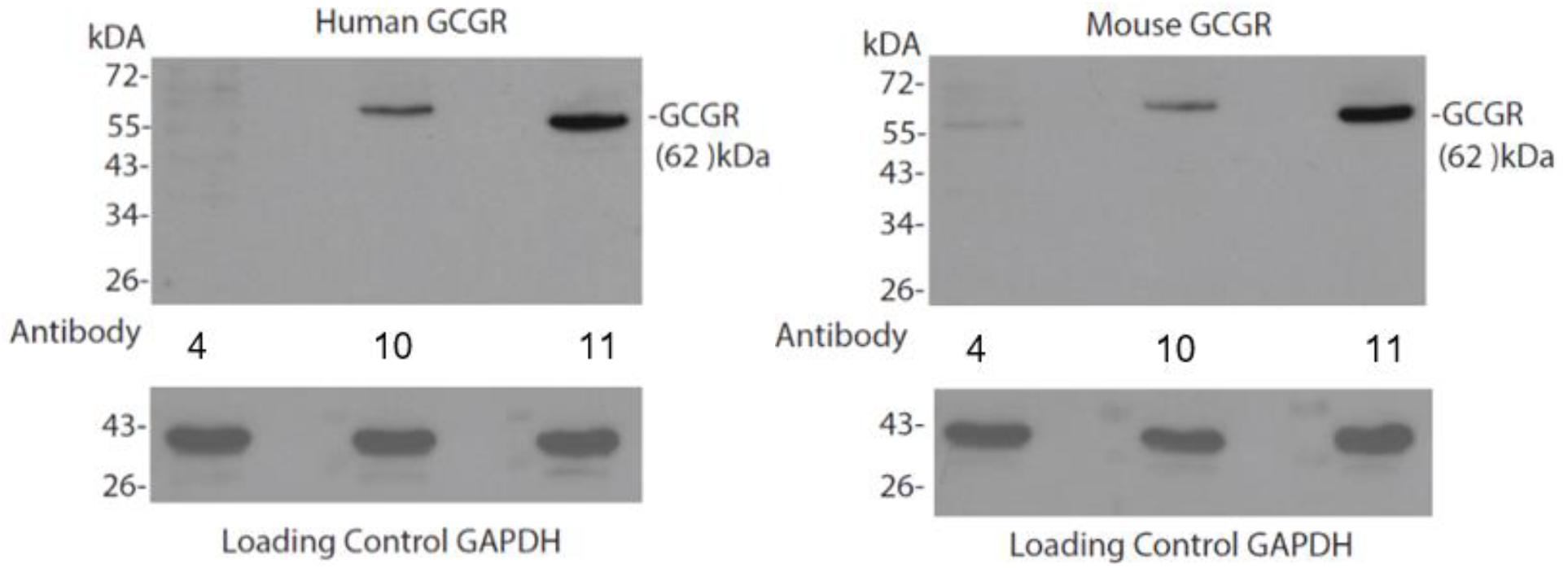
Cropped western blots of transfected cells with either human glucagon receptor (GCGR) cDNA or mouse GCGR cDNA. Western blotting performed with antibody 4, 10 and 11.

To further validate the specificity of antibody no. 11, western blotting was performed on liver tissue from *Gcgr^+/+^* and *Gcgr^−/−^* mice. Western blotting of liver tissue from *Gcgr^+/+^* mice confirmed the specificity of antibody no. 11, detecting a 55kDa band, corresponding to the antibody/GCGR receptor complex (Supplementary figure 3).

### Specific localization of GCGR expression

#### GCGR expression in various mouse tissue using immunohistochemistry

Based on the results obtained from the validation of the twelve commercially available antibodies. Antibody no. 11 was selected for specific localization of GCGR expression in various tissue. Mouse tissue from the following organs were prepared for extensive GCGR immunostaining: heart, pancreas, kidney, liver, gastric mucosa, plexus myentericus, duodenum, ileum, colon, brown and white adipose tissue, and adrenal gland (Figure 5).

**Figure 5.**
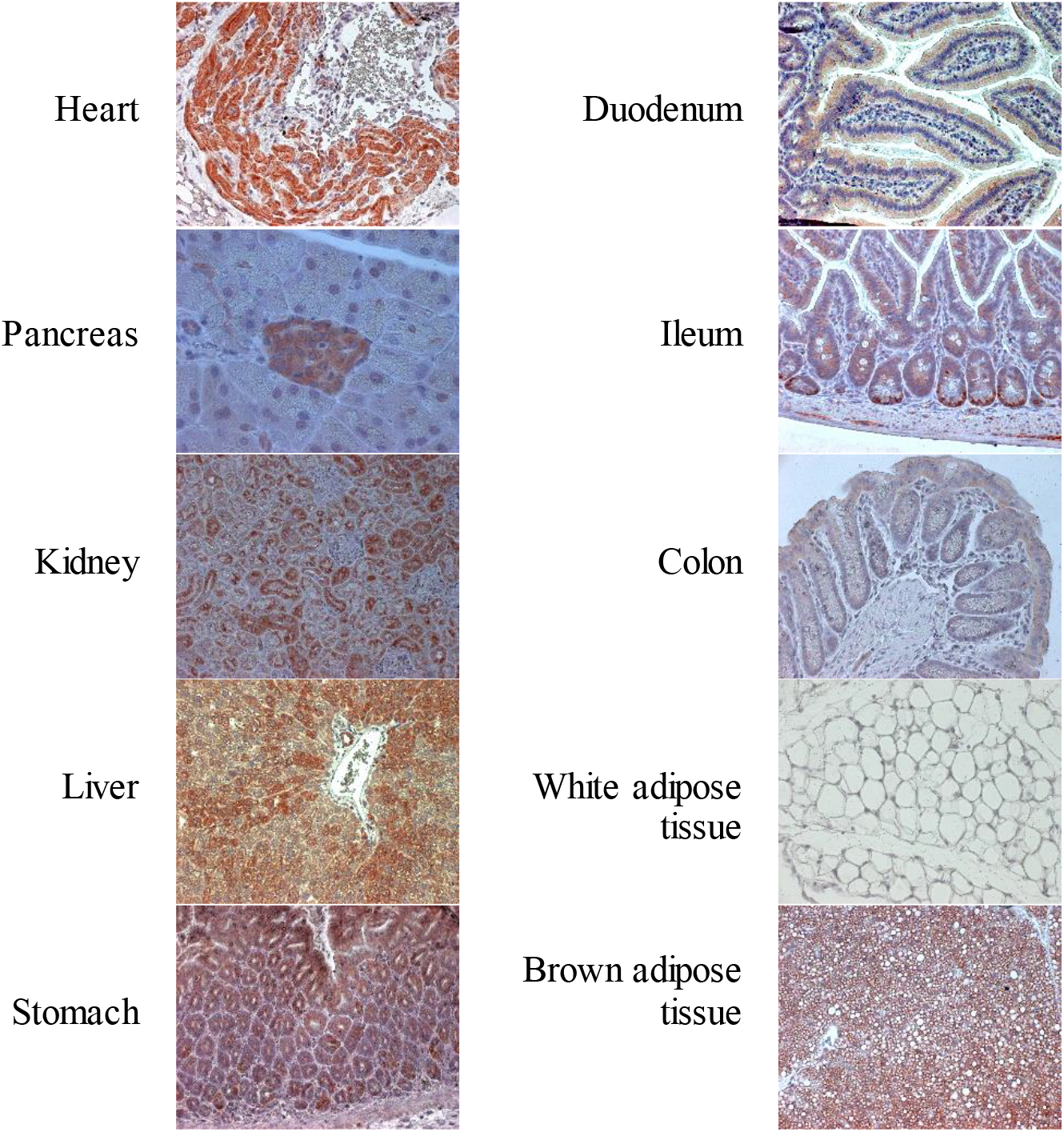
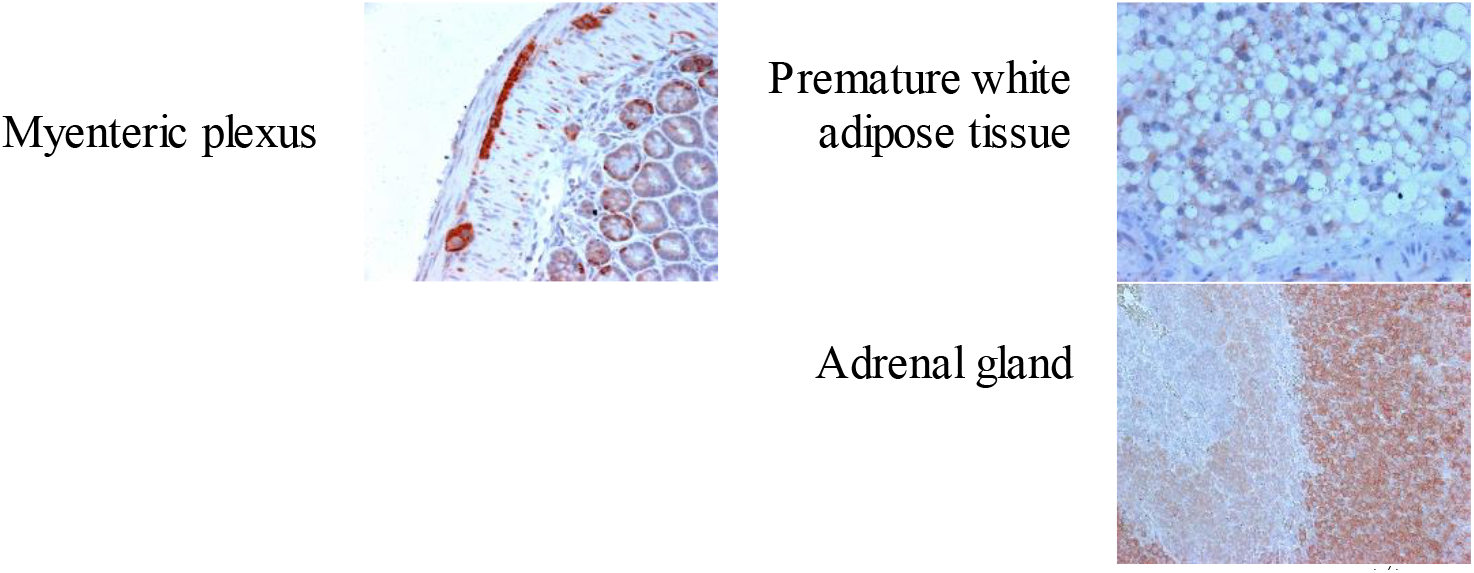
Immunohistochemical staining of various tissues from glucagon receptor wildtype (Gcgr^+/+^) female mice, 8 weeks of age, using the selected antibody no. 11. Immunostainings were found in following tissues: heart, pancreas, kidney, liver, gastric ventricle, nerve fibers, ileum, brown adipose tissue (BAT), premature white adipose tissue (WAT) and the cortex of the adrenal gland. x40, scale bar = 40μm.

The most intense immunostaining reactions were found in heart muscle fibers, kidney tubuli, including both proximal and distal tubules, liver tissue and the islets of pancreas. Glandular cells from stomach and enterocytes in crypts from the ileum were moderately stained and, in some areas, epithelial endocrine-like cells were stained. Further, plexus myentericus nerve fibers were strongly stained for GCGR. Duodenal as well as colonic epithelia were generally negative for GCGR or weakly positive. Concerning white adipose tissue (WAT), the mature fat cells were negatively stained, whereas preadipocytes found in relation to kidney tissue were moderately stained. Finally, thymus cells with adipocyte infiltration were strongly stained for GCGR. In addition, the cortical part of the adrenal gland, but not the medulla was strongly stained for GCGR (Figure 5).

#### GCGR expression in human kidney tissue using immunohistochemistry

Based on the intense immunostaining of kidney tubules cells, human renal tissue from a healthy individual was also examined for localization of GCGR expression using antibody no. 11. Strong immunoreactivity was observed in the distal tubules. Cells within the proximal tubule were negatively stained (Figure 6).

**Figure 6.**
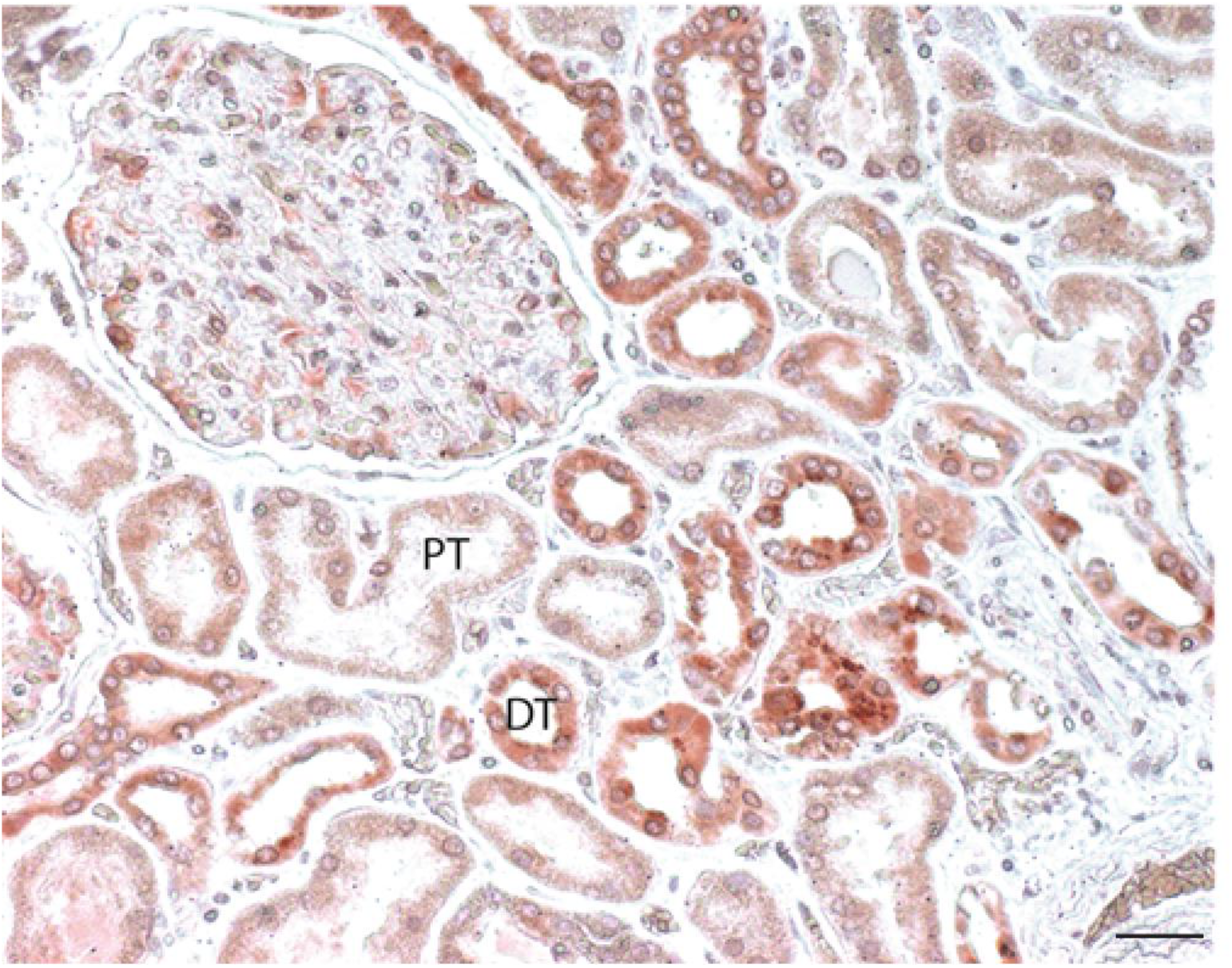
Antibody staining of human renal tissue from autopsy using the selected antibody no. 11. Cells within the proximal tubule (PT) were non-reactive, whereas cells in the distal tubules (DT) showed glucagon receptor (GCGR) immunoreactivity. x100, scale bar = 100 μm

#### Co-staining of pancreatic tissue using immunohistochemistry

GCGR expression in the pancreas is widely debated and we therefore evaluated the cellular location of the GCGR, using antibody no. 11 and co-staining for either glucagon, insulin, or somatostatin. The co-staining with glucagon and antibody no. 11 suggested presence of GCGR in the alpha-cells. Co-staining with insulin and antibody no. 11 also revealed GCGR presence on beta-cells (Figure 7). Further, GCGR expression in the delta-cells was also suggested by co-staining for somatostatin and antibody no. 11.

**Figure 7.**
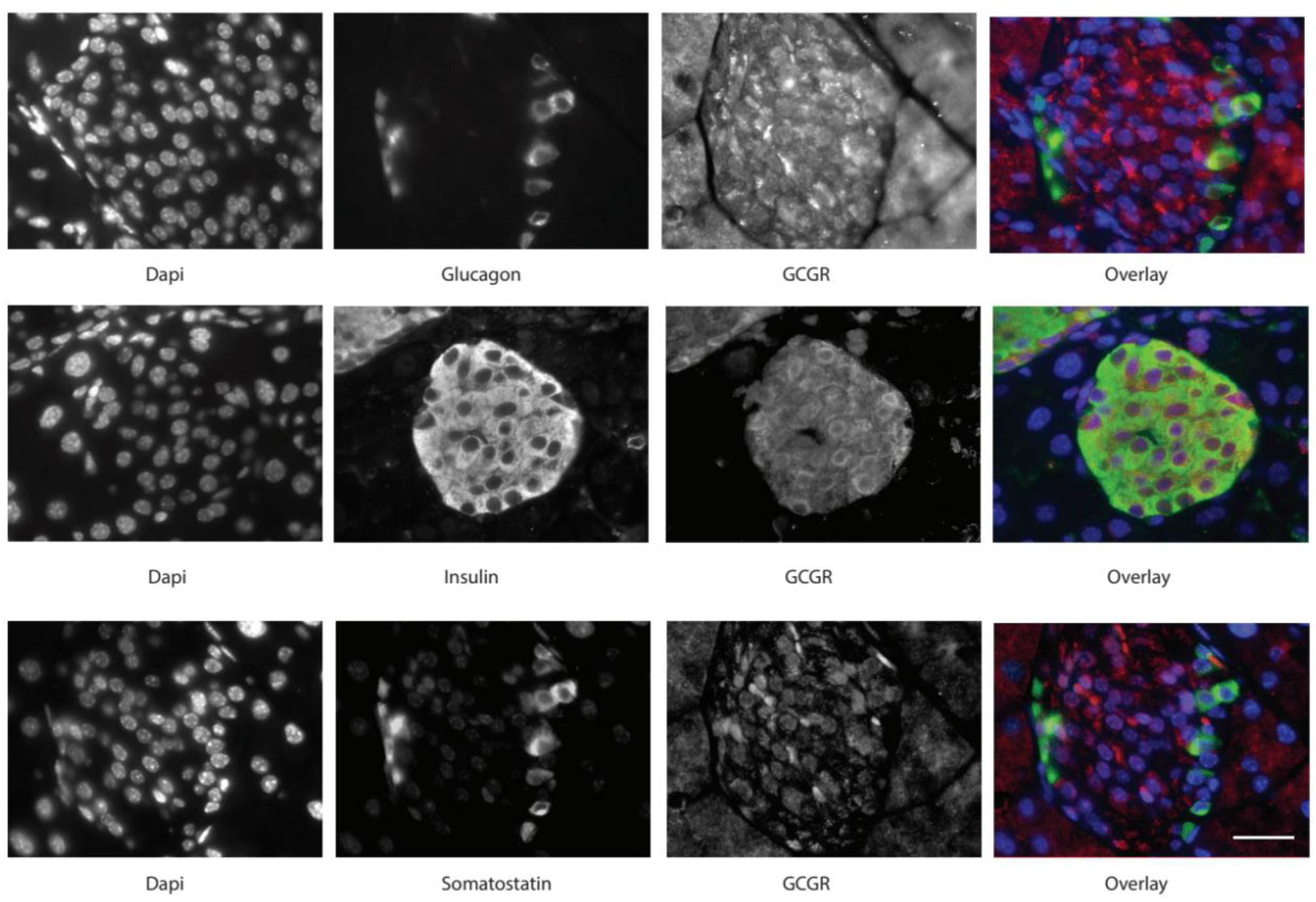
Co-staining of pancreatic mouse tissue, from female mice 8 weeks of age, with antibody no. 11 and either glucagon, insulin, or somatostatin. DAPI staining of nuclei. x85, scale bar = 50 μm.

#### Comparison of glucagon receptor expression by antibody and antibody-independent approaches

Finally, we compared the findings using the GCGR antibody no. 11 to results obtained using an antibody-independent approach: autoradiography. We choose autoradiography as it is a highly sensitive method that depends on ligand-receptor binding, in this case a ^125^I-labelled glucagon molecule. To control for unspecific binding (e.g. to the GLP-1R) we added, in a series of parallel experiments, a 1,000-fold excess of non-labelled glucagon to the ^125^I-Glucagon tracer thereby ensuring imbalanced competition to the GCGR of the non-labelled glucagon molecule allowing us to discriminate between true GCGR binding and unspecific binding.

High density of ^125^I-labeled glucagon grains was observed in hepatocytes whereas in mice receiving both ^125^I-labeled glucagon and an excess of nonradioactive glucagon, a grain density corresponding to background density was observed (Figure 8) supporting the specificity of the result. Similarly, high grain density was observed in the distal and collecting duct cells of the kidney (Figure 8) which was abolished in mice receiving non-labelled glucagon. These findings complement the positive staining of distal and collecting duct cells using antibody no. 11. When compared to both the surrounding exocrine tissue and to the mice who received an excess amount of nonradioactive glucagon, there appeared to be a weak increase in the number of grains in the islets of Langerhans.

**Figure 8.**
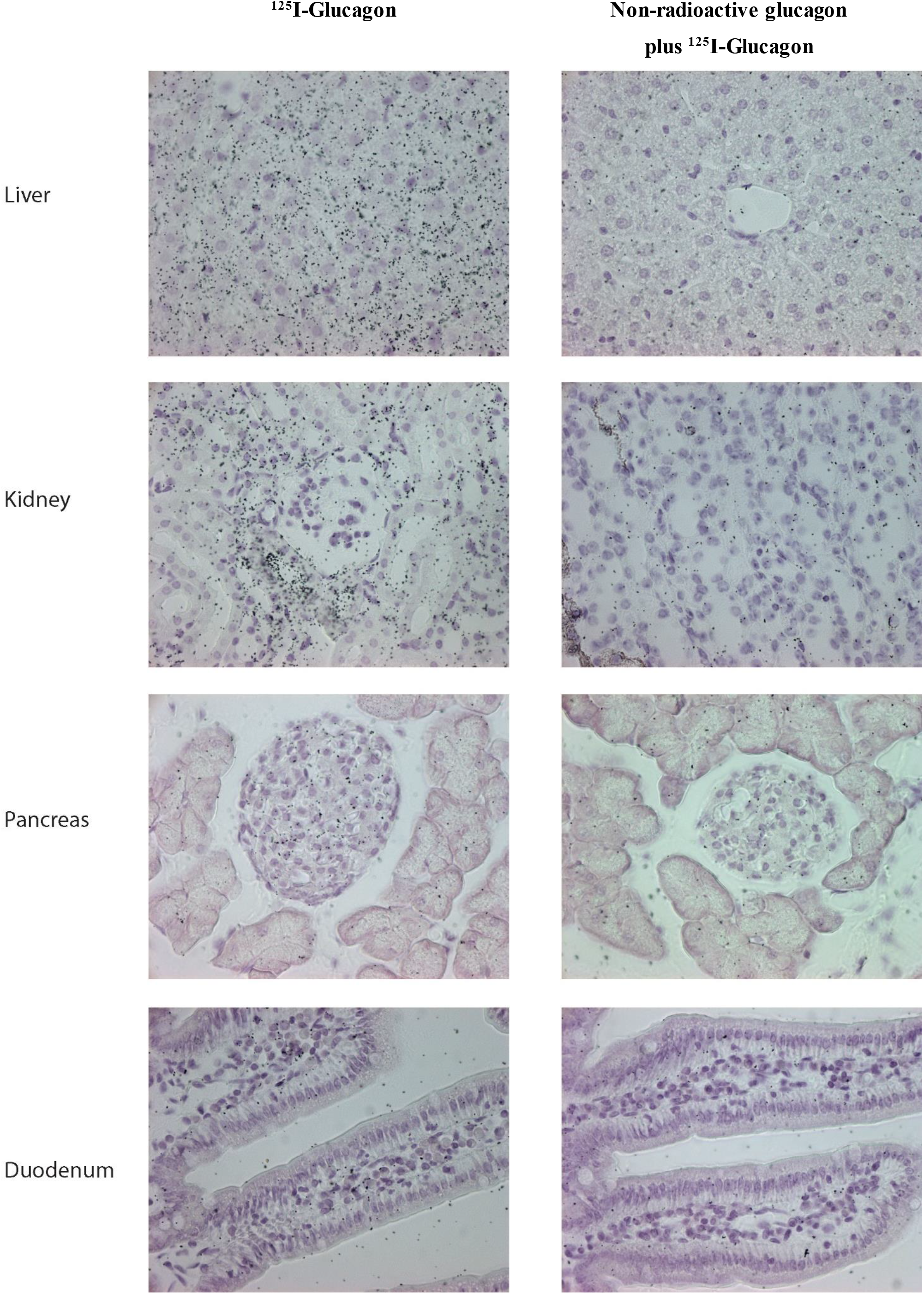
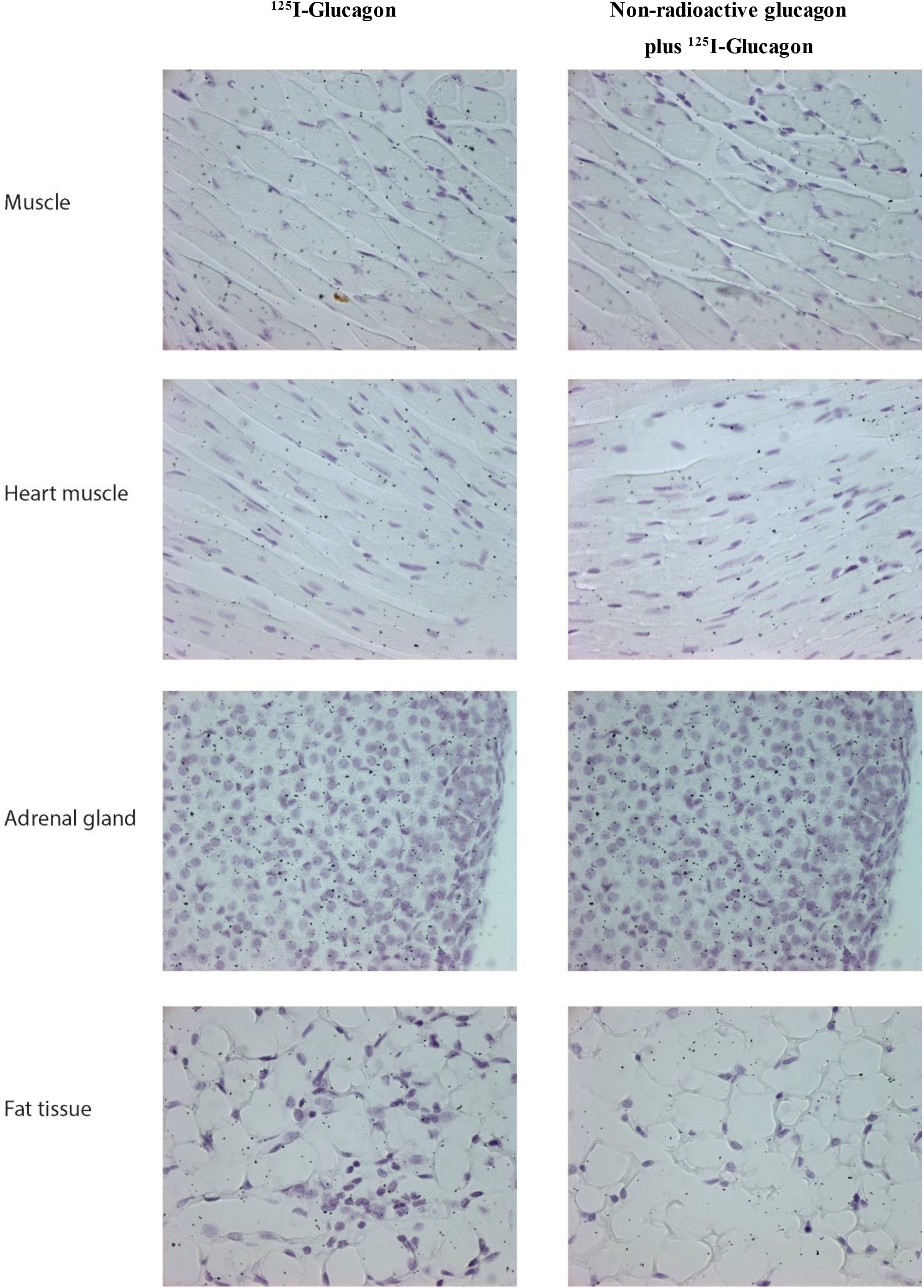
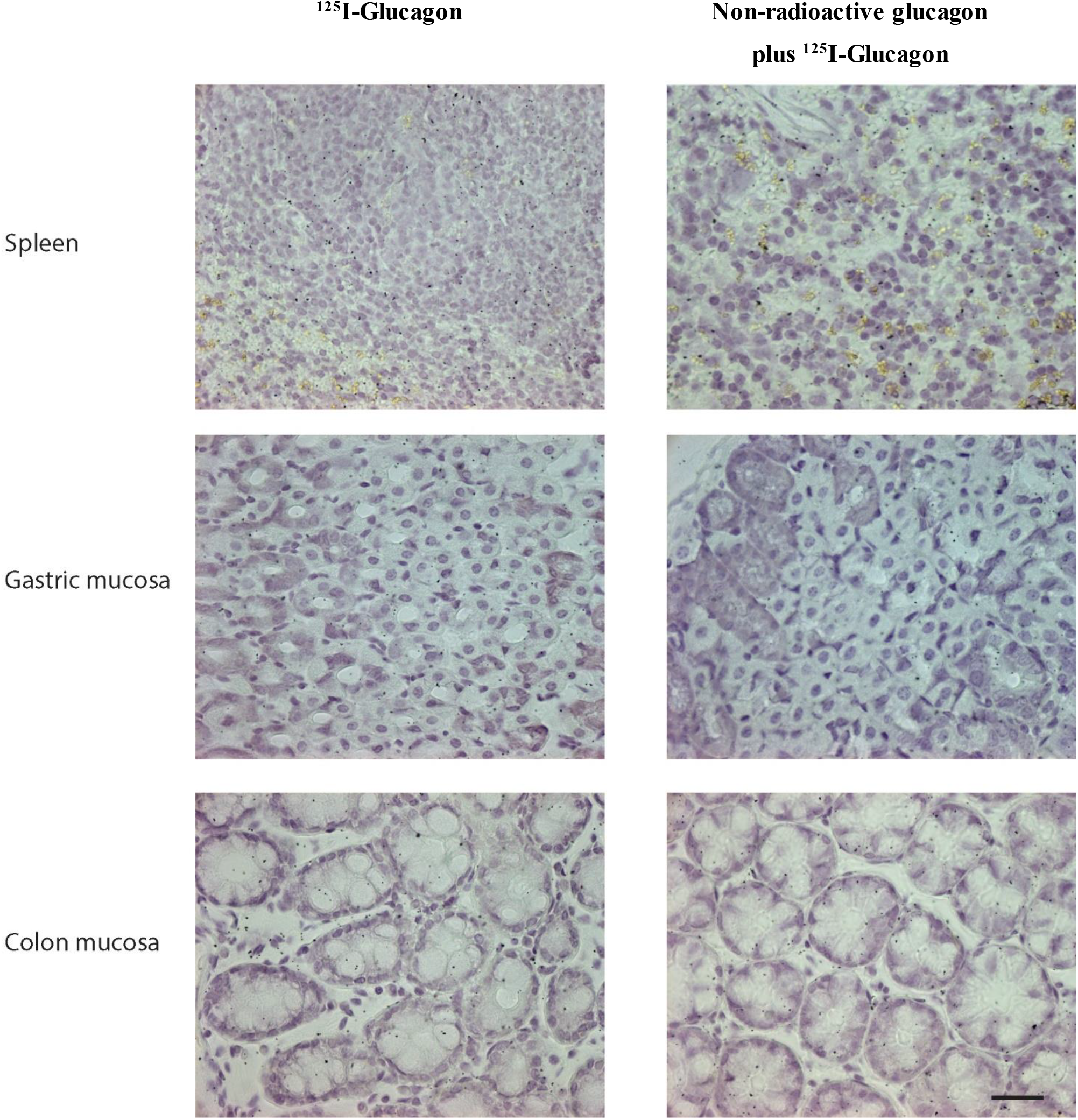
Microphotographs of the liver, kidney, pancreas, duodenum, muscle tissue, heart, adrenal glands, fat tissue, spleen, gastric mucosa, and colon mucosa of female C57BL/6JRj mice, 13 weeks of age, that received an intravenous injection of 3 pmol ^125^I-Glucagon or 5 nmol of non-radioactive glucagon in combination with 3 pmol of ^125^I-Glucagon to test for specific binding. Autoradiography of mouse lever, kidney and pancreas showing high density of grains in the tissue, which was attenuated in mice receiving both labelled and non-labelled glucagon. x120, scale bar = 50 μm.

The positive immunostainings of the stomach, heart, and adrenal glands all suggested GCGR receptor presence within these tissues, however, using autoradiography no grains were present within these tissues. This could indicate lack of receptor expression on the cell surface (Figure 8), perhaps together with an intracellular accumulation of non-functional GCGR protein. However, an inferior sensitivity of the autoradiography approach cannot be excluded. Regarding tissue from the small intestine, muscles, adipose tissue, and spleen neither positive immunostaining nor autoradiografic grains were found (Figure 8).

#### RNA expression of the human GCGR from various tissues and specific cells

To further supplement our findings by antibody and antibody-independent approaches we used RNA-sequencing data of adult human tissues that were made available by the GTEx consortium^25^. We selected tissues with at least ten available profiles sampled from donors who died from natural or violent causes and donors who died unexpectedly with a terminal phase of 1-24 h. This resulted in a total of 22 tissues and 5877 transcriptomic profiles (Supplementary table 2). Varying expression levels of GCGR mRNA were observed with the highest expression (transcripts per million (TPM)) in liver, kidney, and nerve tissue. GCGR mRNA expression in all other analyzed tissue were minimal or not present (Figure 9).

**Figure 9.**
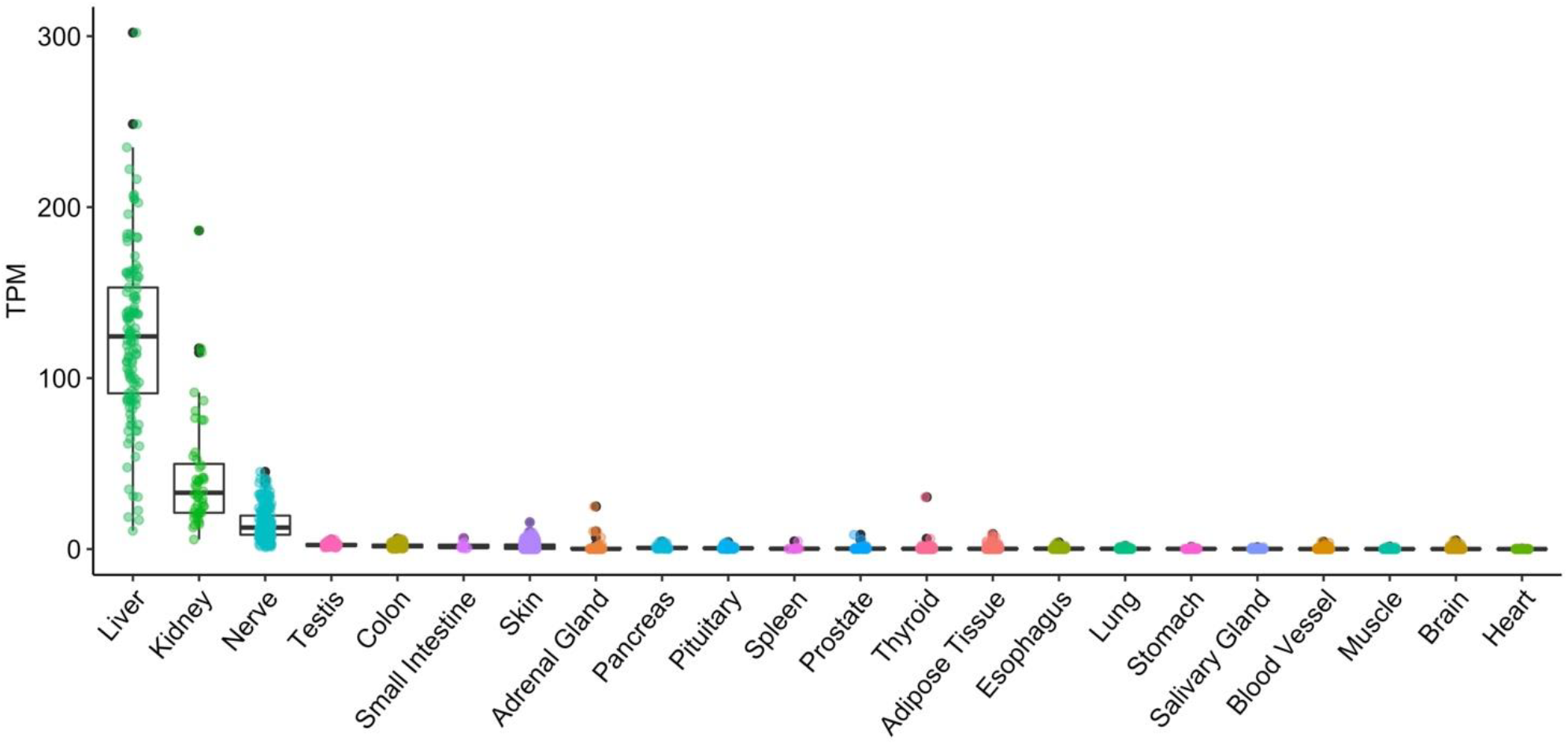
RNA-sequencing data generated by the Genotype-Tissue Expression (GTEx) project^25^ from human tissues displayed in transcript per million (TPM) values for GCGR. GCGR distribution across the dataset is visualized with boxplots, shown as median, 25^th^, and 75^th^ percentiles. Points are displayed as outliers if they are above or below 1.5 times the interquartile range. TPM values of the individual samples are presented with the box plots. The tissues are arranged according to average expression levels from left to right.

To increase our resolution and enable detection of cell specific GCGR mRNA expression, we used 10X Chromium scRNA-sequencing data from MacParland et al., 2018^20^ and Liao et al., 2020^26^ for evaluating single-cell mRNA expression in liver and kidney, respectively. These tissues were selected based on the bulk RNA-sequencing data. The two data sets contain 8444 and 23368 cells, respectively, which passed the quality control restrictions (see the “Material & Methods” section). The analysis and cell type annotations of each cluster are based on the aforementioned publications. The distribution of cell types can be seen in supplementary table 3. For liver tissue, GCGR expression was primarily observed in hepatocytes (Figure 10). Generally, the GCGR expression of the kidney tissue is lower compared to the liver tissue, but the positive cells were primarily observed in collecting and distal tubule cells (Figure 10). Considering scRNA-sequencing being a tertiary approach to support the already confirmed GCGR expression in kidney via IHC, autoradiography, and bulk RNA-sequencing, the 10X data suggest that the expression originates from the collecting and distal tubule cells.

**Figure 10.**
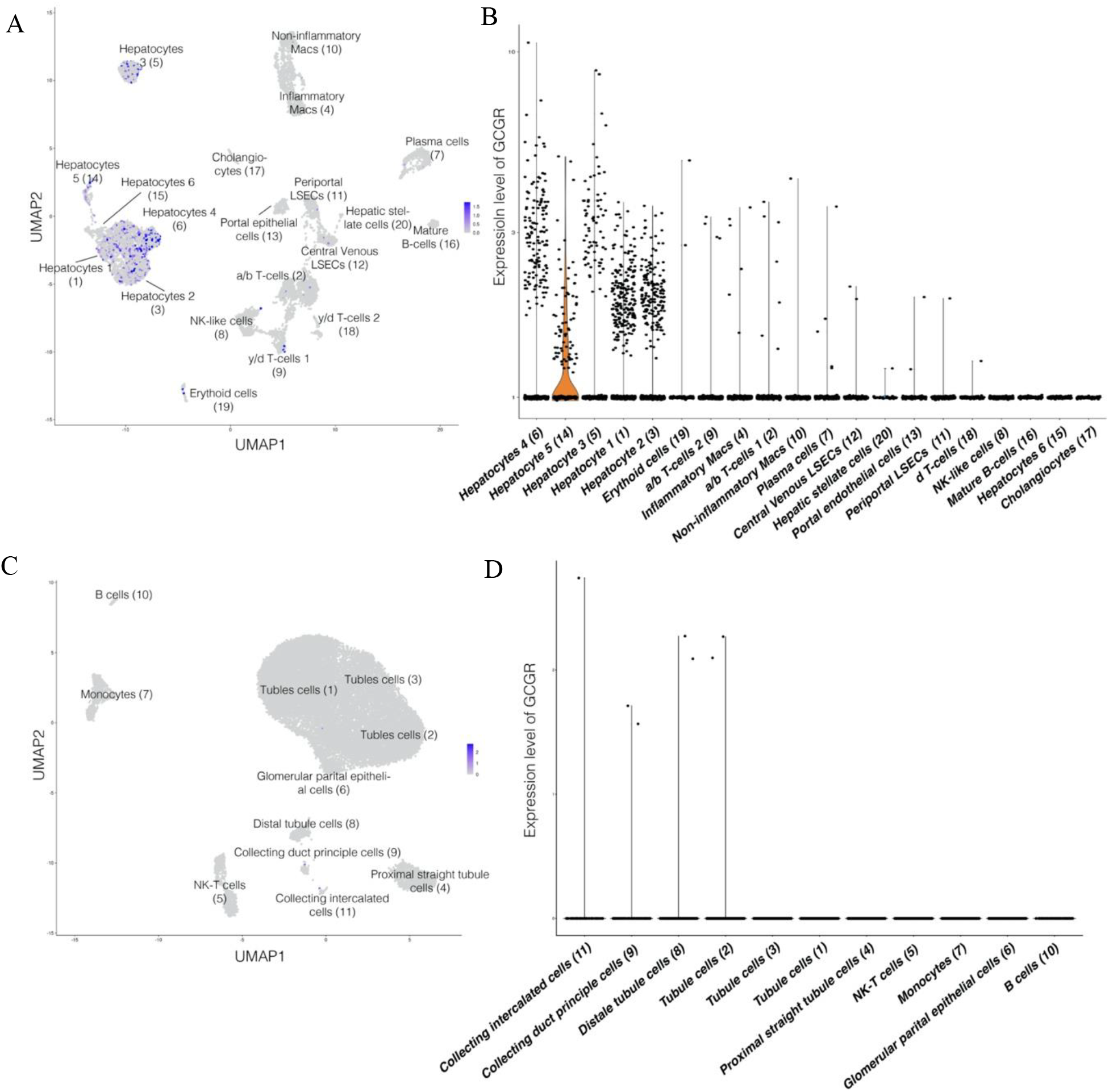
Single-cell RNA-sequencing data of liver and kidney tissue generated by MacParland et al., 2018^20^ and Liao et al., 2020^26^, respectively. A) Feature plot based on a UMAP projection of 8444 hepatic cells with cell type annotations manually annotated by MacParland et al., 2018^20^. GCGR expression is interpretable by the color gradient. B) Violinplot showing the expression of GCGR in the hepatic data set grouped by the annotated cell type arranged according to average expression. C) Featureplot based on a UMAP projection of an integrated data set containing 23368 renal cells derived from three biopsies, showing GCGR expression via the color gradient. Cell type annotations are based on manual annotations by Liao et al., 2020^26^. D) Violinplot showing the expression of GCGR in the renal data set grouped by cell type arranged according to average expression.

#### Overview of glucagon receptor expression by antibody and antibody-independent approaches

## Discussion

Reliable IHC detection of the GCGR depends on high affinity and selectivity of the antibodies used. However, detection of the GCGR expression using IHC, has been hindered by the lack of specific antibodies, limiting the number of valid histological studies. In this report, we demonstrate that only one out of twelve commercially available antibodies provided a specific staining of the GCGR, and based on antibody-independent approaches, we were able to confirm the selectivity of this antibody. This confirms the difficulty regarding IHC specificity in attempts to localize GPCR – an issue which has been addressed comprehensively^35–37^. A similar pattern of unspecific reactions has been observed in wild-type and knockout mice, which might suggest that antibodies tend to bind to cells by antigen-independent mechanisms thus giving ambiguous results. In general, GPCR proteins are expressed in low quantities, making them hard to detect using antibody approaches. The low expression likewise poses a challenge when investigating the expression of GPCRs through RNA-sequencing based approaches. GPCR typically have high degrees of homology and antibodies may therefore easily recognize other subtypes within the same family. This could explain the unspecific reaction patterns of the antibodies observed within this study. This phenomenon was previously documented in a study of adrenoceptors, for which antibodies directed against either β2- or β3-adrenoceptors also recognized bands on immunoblots from cells expressing any of the nine adrenoceptors subtypes^38^, illustrating their promiscuous nature.

The antibodies used in this study are all polyclonal. It is commonly assumed that polyclonal antibodies are more likely to cross-react with other proteins than monoclonal antibodies, suggesting that monoclonal antibodies should be preferred. However, polyclonal antibodies have, in diagnostic applications, previously been demonstrated to be advantageous due to their multi-epitope binding properties^39,40^, which has been shown to increase sensitivity when used for detecting proteins that are present in low quantities^41^. In addition, some of the clones may have a higher binding affinity than the average affinities of the monoclonal antibodies, a feature that is exploited in radioimmunoassays, where the high-affinity clones govern the sensitivity of the assay.

Besides specificity, other factors have been demonstrated to influence the binding of antibodies to their targets, such as methods of cell/tissue fixation, tissue processing, and detection methods^42^. Especially fixation has a major impact on affinity and selectivity of antibodies since fixation changes the chemical property of the tissue, which alters the three-dimensional protein conformation by cross-linking^43^.

Given the many disadvantages of IHC, avoiding IHC approaches may be desirable. In an attempt to localize the GCGR, many studies have resorted to methods such as RT-PCR^1,4^, ribonuclease protection assay^3^, or northern blotting analysis^5^ for estimation of GCGR mRNA levels. The expression of GCGR mRNA was found in various tissues, such as the liver^1–3^, kidney^1–3^, heart^1–3^, adipose tissue^1–3^, spleen^3^, pancreatic islets^3^, stomach^1,3^, small intestine^1,3^, adrenal glands^1–3^, and skeletal muscle^3^. It has, however, been suggested that the GCGR mRNA and protein expression at the cell surface do not necessarily correlate, since the major regulation of the latter *in vivo* takes place at a posttranscriptional level^6^. It has been established that the amount of functional GCGR protein is lower than the GCGR mRNA, rendering the use of GCGR mRNA expression alone unsuitable for tracing of receptor protein expression on the cell surface. Another important issue with posttranscriptional methods is their inability to identify specific cellular localization of the GCGR. Indeed, measurement of the protein expression would be necessary for exact GCGR expression quantity and specific localization, as evident for analysis of receptor localization in nerve terminals.

Based on our results, the present findings suggest that additional, non-antibody-based validation approaches should be applied together with the antibody-based approaches to make up for possible cross-reactivity and selectivity. To confirm the antibody’s diagnostic sensitivity and specificity even further, reproducibility using other lots of the same antibody ought to be applied when using polyclonal antibodies.

### Specific localization of GCGR expression

Since glucagon primarily acts on liver GCGR to increase blood glucose by promoting glycogenolysis and gluconeogenesis^44^, most studies focus on the GCGR function in the liver. However, GCGR has also been detected in many other tissues, including the kidney, heart and adipose tissue^2,45^. In this study, we searched for the specific location of the GCGR using an antibody-based approach and two non-antibody-based approaches: autoradiography and data analysis from RNA-sequencing data. Using IHC, various mouse tissues were examined for GCGR expression. The most intense immunostainings were observed in the liver, heart muscle fibers, kidney tubuli (including both proximal and distal tubuli), the islets of pancreas, plexus myentericus nerve fibers, and preadipocytes found in relation to kidney tissue. The specific GCGR localization in some of these different tissues has been the subject of vivid discussions. Profound disagreements regarding direct glucagon action and GCGR localization are apparent.

In the pancreas, the presence of the GCGR has previously been demonstrated, by measuring GCGR mRNA^46^. Controversies exist whether the insulinogenic effect of glucagon is mediated directly via the GCGR on beta-cells. However, increasing concentrations of glucose have previously been demonstrated to upregulate the GCGR in beta-cells prompting GCGR action necessary to maintain beta-cell glucose competence^47^. By immunohistochemistry activity the GCGR, besides in the beta-cells, has also been identified in rodent pancreatic alpha- and delta-cells^33^, suggesting that these cells may be influenced by glucagon in a paracrine manner and that glucagon might even bind to receptors on the alpha-cell, in an autocrine manner^48^. Such notions are in agreement with the results found in this study: we demonstrated the presence of the GCGR in the alpha- and beta- cells, using a co-staining method with antibody no. 11 and either glucagon or insulin.

In the adipose tissue, it has been suggested that glucagon serves a direct role, by increasing blood flow, stimulating lipolysis, increasing glucose uptake and increasing oxygen consumption in white- and brown-adipose tissue^49–52^. This indicates that the GCGR is most likely expressed on these cells. A specific localization of the GCGR has also been identified in both white- and brown- adipose tissue^53,54^, predominantly based on the identification of GCGR mRNA transcripts^2,46^. These studies generally observed a relative abundance of GCGR mRNA, which was highest in brown adipose tissue followed by white adipose tissue. In the present study we found a moderate staining of preadipocytes while no white adipose tissue was stained, using antibody no. 11. This could be due to different methodologies where the PCR detects the GCGR mRNA transcripts. Similar to our findings, another study observed poor immunoblots of adipose tissues, when using a monoclonal GCGR antibody^27^. These observations could in fact be due to low expression levels of protein GCGR. However, we were also unable to identify mRNA GCGR transcript from the RNA sequencing data of adult human tissues processed within this study, where no expression levels of GCGR mRNA was observed in adipose tissue. These findings are in alignment with previous finding, demonstrating that glucagon does not stimulates lipolysis in adipose tissue^55^, which may suggest that GCGR is not present in white adipose tissue.

The adipose tissue is not the only location for questionable GCGR localization. We located the GCGR in the heart muscle fibers, using an IHC approach, but we were unable to detect the GCGR using the antibody-independent approach, autoradiography. Furthermore, it was not possible to identify GCGR mRNA transcripts from the RNA-sequencing data of adult human heart tissue. In previous studies, GCGR mRNA transcripts have been detected in heart tissue of both rats^2^ and mice^56^, however, not in humans^57^. Therefore, the common understanding of glucagon action and specific GCGR localization in the heart remains limited. GCGR mRNA has been detected from both right and left atria and ventricle of the adult mouse heart, using RT-PCR^56^, where others have found GCGR distribution considerably higher in ventricular than in atrial myocardium by using western blotting^58^. This demonstrates the degree of difficulty in detecting receptors that are weakly expressed.

We found GCGR in nervous tissue with antibody no. 11 and we also found mRNA transcripts from the RNA-sequencing data of adult human tissues. Demonstrating strong evidence for presence of the GCGR in plexus myentericus nerve fibres and tibial nerves, respectively. This could suggest a new unidentified localization of the GCGR, since there, to our knowledge, is no literature describing this. The location of GCGR in these tissues would point to the existence of hitherto unrecognized glucagon actions and should be investigated further.

The involvement of GCGR in kidney function has recently emerged in the scientific field since glucagon has been shown to cause glomerular hyperfiltration^59^, and to acutely increase urinary excretion of urea, sodium, and potassium^60^. This raises new questions concerning the role of glucagon in kidney function. In this study, we observe specific GCGR localization in the distal tubule in human tissue, using a selected antibody. The presence of GCGR in renal tissue was supported by autoradiography, where ^125^I-labeled glucagon grains were observed in distal and collecting duct cells of the kidney. RNA-sequencing data confirmed GCGR presence in liver and kidney tissue while the scRNA-sequencing data clearly indicates that the GCGR is expressed in most types of hepatocytes, while the much lower expression of GCGR in kidney seems to be restricted to collecting and distal tubules cells. The ambiguity of the kidney scRNA-sequencing data might be due to either limited coverage, insufficient sequencing depth, or simply very low GCGR expression. The expression of GCGR in the distal part of the kidney supports a physiological role for glucagon in this target organ. Several studies have suggested that GCGR activation influences the transport of fluid and solutes in the distal tubule and collecting duct in rats and humans^61–63^. Our data on GCGR localization in the kidney support these findings and may help explain its effects in the distal tubule. Our findings are in agreement with previous findings of GCGR in the entire distal nephron but not in the proximal tubule^32,64^. However, glucagon has also been demonstrated to exert vascular effects in the kidney and to induce renal vasodilation in different animal models^65,66^. In humans, glucagon has been suggested to reduce tubular protein reabsorption^59^. Our data on GCGR localization in normal kidney tissue do not directly explain these effects. The precise localization of the GCGR and its sites of action in the functional nephron are critical for understanding the interactions between the kidney and the endocrine system in relation to fluid volume homeostasis, blood pressure control, biochemical and metabolic regulation, and deserves further investigation.

In summary, the evaluation of twelve commercially available GCGR antibodies resulted in the conclusion that GCGR localization remains challenging due to the lack of suitable commercially available GCGR antibodies. However, based on extensive validation of the IHC approach and the use of supplementary approaches, the localization of the GCGR could be estimated. The present study identifies GCGR localization sites in the liver, pancreas, preadipocytes, heart muscle fibers and kidney. The specific GCGR localization in the kidney should be explored further as it indicates the existence of a potential, hitherto unrecognized, pancreas-renal interplay.

## Acknowledgments

A sincere thanks to Maureen J. Charron, Departments of Biochemistry, Obstetrics and Gynecology and Women’s Health, and Medicine, Albert Einstein College of Medicine, New York, New York for providing glucagon receptor knockout mice.

## Author Contributions

J.J.H., R.A., and N.J.W.A. conceived and planned the experiments. A.B.B., C.D.J., S.A.S.K., K.D.G., J.B.C., M.W.S., C.Ø., and R.A. carried out the experiments. C.D.J., J.B.C., and N.J.W.A. carried out data interpretation using database available RNA-sequencing and scRNA-sequencing data. A.B.B., S.A.S.K., K.D.G., M.W.S., C.Ø., R.A., and N.J.W.A. contributed to the interpretation of the results. M.H. and E.P., collected and provided healthy human kidney biopsies. R.S and L.L.G. performed and provided liver biopsies from patients with NASH. A.B.B., C.D.J., J.B.C., S.A.S.K., K.D.G., J.J.H., R.A., and N.J.W.A. drafted manuscript. N.J.W.A. is responsible for project administration. All authors provided critical feedback and helped shape the research, analysis and manuscript.

## Data Availability

The data sets generated are not publicly available but can be made available from the corresponding author upon reasonable request. Script for data processing of single-cell and bulk RNA-sequencing is available at GitHub (https://github.com/nicwin98/GCGR_Expression).

