## Supplementary material for "Evaluation of Commercially Available Glucagon Receptor Antibodies and Glucagon Receptor Expression": Supplemantary Material

### Supplementary table 1

**Supplementary table 1.** *Sequencing of plasmids used for glucagon receptor (GCGR) expression levels investigation.*

| |
|---|
| To ensure the right transcript was used, plasmids from both human and mouse were sequenced.<br>GCGR transcript from both the human and mouse plasmids was found, demonstrated below. |
| >H1<br>TNNNGNAATGGGCGGTAGGCGTGTACGGTGGGAGGTCTATATAAGCAGAGCTCGTTTAGT<br>GAACCGTCAGAATTTTGTAATACGACTCACTATAGGGCGGCCGGGAATTCGTCGACTGGA<br>TCCGGTACCGAGGAGATCTGCCGCCGCGATCGCCATGCCCCCCTGCCAGCCACAGCGACC<br>CCTGCTGCTGTTGCTGCTGCTGCTGGCCTGCCAGCCACAGGTCCCCTCCGCTCAGGTGAT<br>GGACTTCCTGTTTGAGAAGTGGAAGCTCTACGGTGACCAGTGTCACCACAACCTGAGCCT<br>GCTGCCCCCTCCCACGGAGCTGGTGTGCAACAGAACCTTCGACAAGTATTCCTGCTGGCC<br>GGACACCCCCGCCAATACCACGGCCAACATCTCCTGCCCCTGGTACCTGCCTTGGCACCA<br>CAAAGTGCAACACCGCTTCGTGTTCAAGAGATGCGGGCCCGACGGTCAGTGGGTGCGTGG<br>ACCCCGGGGGCAGCCTTGGCGTGATGCCTCCCAGTGCCAGATGGATGGCGAGGAGATTGA<br>GGTCCAGAAGGAGGTGGCCAAGATGTACAGCAGCTTCCAGGTGATGTACACAGTGGGCTA<br>CAGCCTGTCCCTGGGGGCCCTGCTCCTCGCCTTGGCCATCCTGGGGGGCCTCAGCAAGCT<br>GCACTGCACCCGCAATGCCATCCACGCGAATCTGTTTGCGTCCTTCGTGCTGAAAGCCAG<br>CTCCGTGCTGGTCATTGATGGGCTGCTCAGGACCCGCTACAGCCAGAAAATTGGCGACGA<br>CCTCAGTGTCAGCACCTGGCTCAGTGATGGAGCGGTGGCTGGCTGCCGTGTGGCCGCGGT<br>GTTCATGCAATATGGCATCGTGGCCAACTACTGCTGGCTGCTGGTGGAGGGCCTGTACCT<br>GCACANCCTGCTGGGCCTGGCCACCCTCCCCGAGAGGAGCTTCTTCNCCTCTACTGGGCN<br>TCGGCTGGNTGCCCCNTNCTGTNN |
| >H2<br>NGAGANGCCTGGGGAGGGGTCACAGGGATGCCACCCGGGATCTGTTCAGGAAACAGCTAT<br>GACCGCGGCCGGCCGTTTAAACCTTATCGTCGTCATCCTTGTAATCCAGGATATCATTTG<br>CTGCCAGATCCTCTTCTGAGATGAGTTTCTGCTCGAGCGGCCGCGTACGCGTGAAGGGGC<br>TCTCAGCCAATCTAGGGAGGCCACCAGCCAAGGGGGTCTCCGCAGATGAATCCTGGCTGC<br>CACCACCCCTCCCAAACTGCAGCTCCTTGCTGGGAGGGCCGTGGCCGGGCGAAGATGAGG<br>CCCTGTGGTTGCTGGTGTTCCGCTCCTCCCATAGCACTTTGCCCAGGCGCCAGCGGTGCC<br>AACGCCGCCGCAGCTCCGACTGCACCTCCTTGTTGAGGAAGCAGTAGAGGACAGCCACCA<br>GCAGGCCCTGGAAGGAGCTGAGGAAGAGGTCGAAGAAGAGCTTGGCGGAGCGCAGGGTGC<br>CCTGGGCGTGCTCGTCCGTCACGAAGGCGAAGACCACTTCGTGGACGCCCAGCAGAGGGA<br>TGAGGGTCAGCGTGGACTTGGCCAGCCGGAACTTGTAGTCTGTGTGGTGCATCTGCCGTG<br>CCCGCAGCTTGGCCACGAGCAGCTGAACGATGCGGACGAAGATGAAGAAGTTGATCAGGA<br>TGGCCAGGAAGACGGGGAACCGCAGGATCCACCAGAAGCCCATGTTGTCATTGCTGGTCC<br>AGCACTGGACGTTCTCGAACAGACACTTGACCACTGCCCAGGGGACGACGAACAGCATGG<br>GGGCACCCCAGCCGATGCCCAGGTAGAGGCTGAAGAAGCTCCTCTCGGGGAGGGTGGCCA<br>GGCCCAGCAGGTTGTGCAGGTACAGGCCCTCCACCAGCAGCCAGCAGTAGTTGGCCACGA<br>TGCCATATTGCATGAACACCGCGGCCNACGGCAGCCAGCCNCCGCTCNN |
| >M1 is M2<br>NAGGAGAGGCCTGGGGAGGGGTCACAGGGATGCCACCCGGGATCTGTTCAGGAAACAGCT<br>ATGACCGCGGCCGGCCGTTTAAACCTTATCGTCGTCATCCTTGTAATCCAGGATATCATT<br>TGCTGCCAGATCCTTCTTCTGAGATGAGTTTCTGCTCGAGCGGCCGCGTACGCGTGGTGG<br>GGCTGTCAGCCAACCTTGGGAGACTACTGGCCAGCGAGGTCTCCATAGAGGGCACACAGC |

| |
|---|
| CAGTCCCACTGCTGCTGCCTGCACTCATAAGCTGAAGTTTCTCACAGGGATCACCATGAC<br>AAGGCCCTGCTGGGGCCATGTGGCTGCCATGGCTGCTGGCCAACCTTTCCTCCTGAAGAG<br>CTTTGCCCTTCTTGCCATTGCCTCCAACGCCGCATCAGCTCTGCCTGCACCTCCTTGTTG<br>AGGAAACAGTAGAGAACAGCCACCAGCAGACCCTGGAAGGAGCTGAGGAACAGGTCAAAA<br>AAGAGCTTGGTGGAGCGCAGGGTGCCTTGGGCATGCTCGTCAGTCACAAAGGCAAAGACC<br>ACCTCGTGGACCCCCAGCAGAGGGATGAGGGTCAGCGTGGACCTGGCCAGCCGGAACTTA<br>TAGTCAGCATAGTGCATCTGATGGGCACGCAGCTTGGCCACAAGAAGGTGAATGATGTGG<br>ACAAAGATGAAAAAATTGATCAGTAAGGCCAGGAAGACAGGAATACGCAGGATCCACCAG<br>AATCCCATGTTGTCATTGCTGGTCCAGCACTGAACATTCTCAAACAGACACTTGACCACC<br>ACCCCAGGGGATGACAAACAGCAGGGGCGCACCCCAGCCAATGCCCAGGTAGAGGGAAAA<br>GAAGCTCCTCTCAGAGAAGGTGGCAAGGCTCAGCAGGCTGTACAGGTACACGCCTCTANA<br>GCACCAGCATAGTTGGCTATGATGCCGTACTGCATGATCANTGTGGCNNTCTGCAGCCGG<br>CNTCGCCCGTNCTGAGC |
| >M2=M1<br>TNNNGNAATGGGCGGTAGGCGTGTACGGTGGGAGGTCTATATAAGCAGAGCTCGTTTAGT<br>GAACCGTCAGAATTTTGTAATACGACTCACTATAGGGCGGCCGGGAATTCGTCGACTGGA<br>TCCGGTACCGAGGAGATCTGCCGCCGCGATCGCCATGCCCCTCACCCAGCTCCACTGTCC<br>CCACCTGCTGCTGCTGCTGTTGGTGCTGTCATGTCTGCCAGAGGCACCCTCTGCCCAGGT<br>AATGGACTTTTTGTTTGAGAAGTGGAAGCTCTATAGTGACCAATGCCACCACAACCTAAG<br>CCTGCTGCCCCCACCTACTGAGCTGGTCTGTAACAGAACCTTCGACAAGTACTCCTGCTG<br>GCCTGACACCCCTCCCAACACCACTGCCAACATTTCCTGCCCCTGGTACCTACCTTGGTA<br>CCACAAAGTGCAGCACCGCCTAGTGTTCAAGAGGTGTGGCCCGATGGGCAGTGGGTTCG<br>AGGGCCACGGGGGCAGCCGTGGCGCAACGCCTCCCAATGTCAGTTGGATGATGAAGAGAT<br>CGAGGTCCAGAAGGGGGTGGCCAAGATGTATAGCAGCCAGCAGGTGATGTACACCGTGGG<br>CTACAGTCTGTCCCTGGGGGCCTTGCTCCTTGCGCTGGTCATCCTGCTGGGCCTCAGGAA<br>GCTGCACTGCACCCGAAACTACATCCATGGGAACCTGTTTGCGTCCTTTGTGCTCAAGGC<br>TGGCTCTGTGTTGGTCATCGATTGGCTGCTGAAGACACGGTACAGCCAGAAGATTGGCGA<br>TGACCTCAGTGTGAGCGTCTGGCTCAGTGACGGGGCGATGGCCGGCTGCAGAGTGGCCAC<br>AGTGATCATGCAGTACGGCATCATAGCCAACTATTGCTGGTTGCTGGTAGAGGGCGTGTA<br>CCTGTACAGCCTGCTGAGCCTTGCCNCCTTCTCTGAGAGGAGCTTCTTTTCCTCTACTGG<br>GCATTGGCTGGGTGCGCCCTGCTGTTGNATCCCNN |

**Supplementary figure 1**

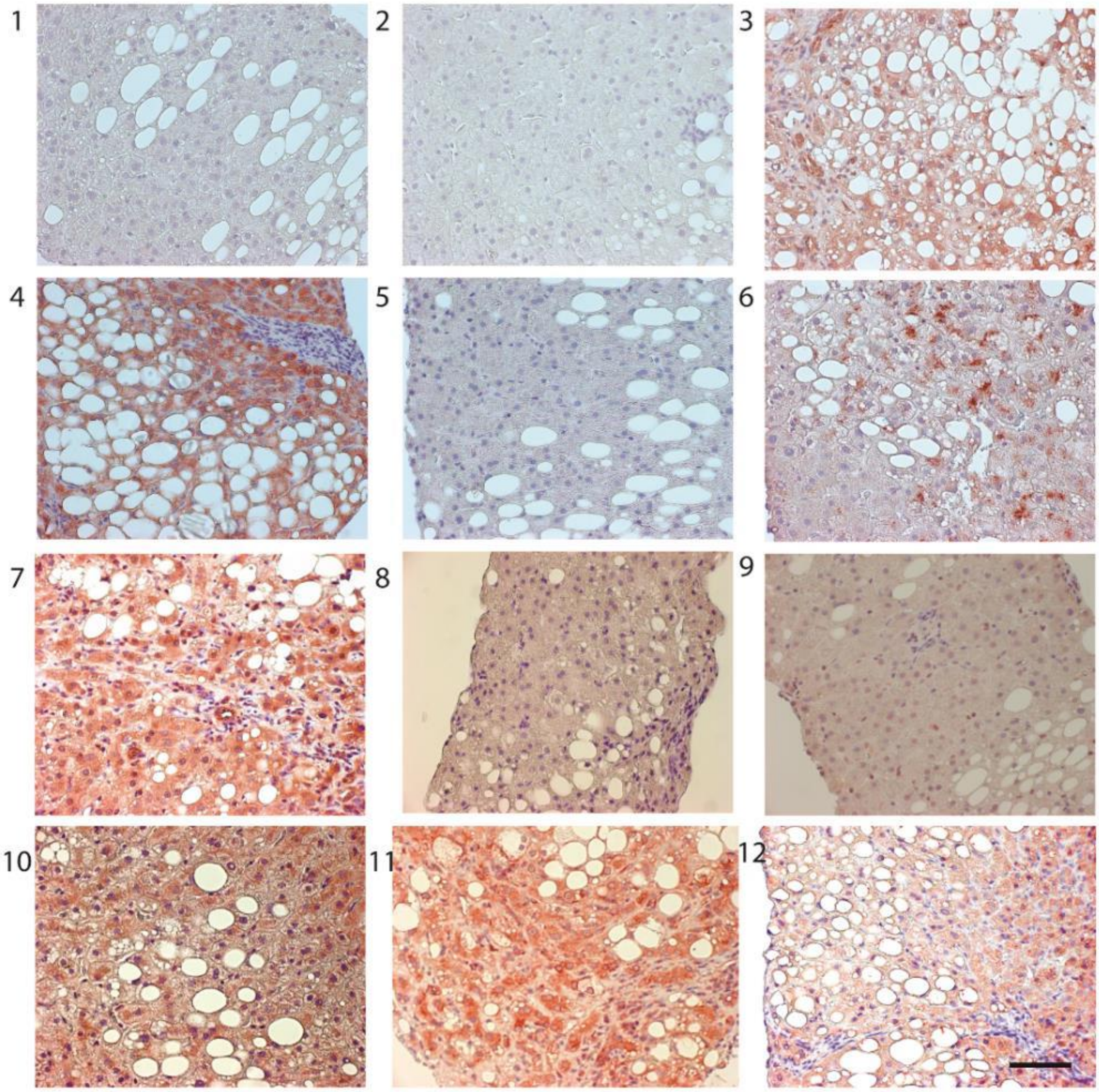


**Supplementary figure 1**. *Paraffin-embedded liver biopsies from humans with non-alcoholic steatohepatitis (NASH) (n=3) stained using the twelve glucagon receptor (GCGR) antibodies. Antibodies no. 3, 4, 6, 7, 10, 11 and 12 showed varying staining intensity. x60, scale bar = 100 µm.*

**Supplementary figure 2**

Antibody no. 4 10 11

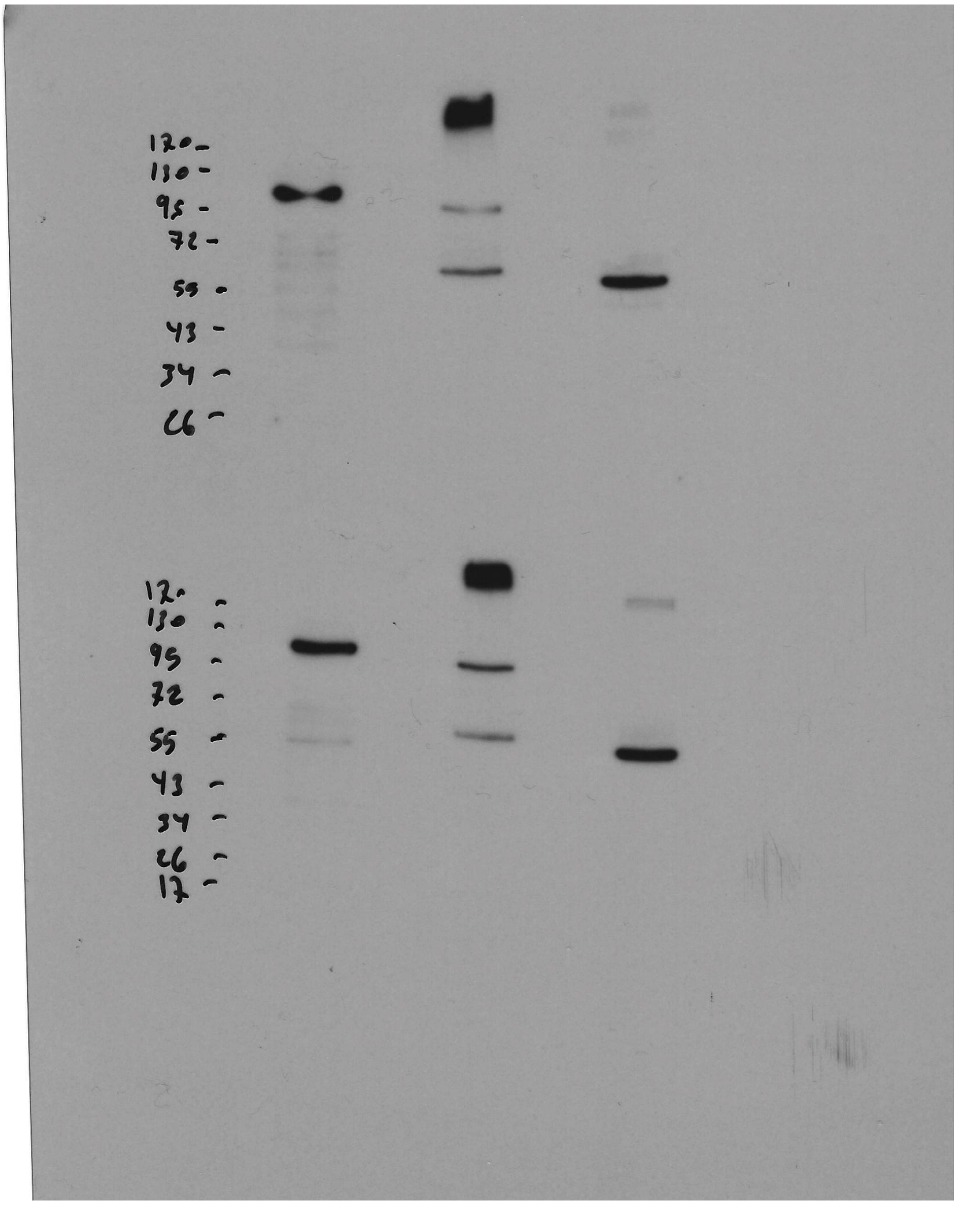

Human GCGR

Mouse GCGR

**Supplementary figure 2.** *Uncropped protein immunoblot. 293-vnR cells transfected with human glucagon receptor (GCGR) and mouse GCGR. Western blotting performed with antibody 4, 10 and 11.*

**Supplementary figure 3**

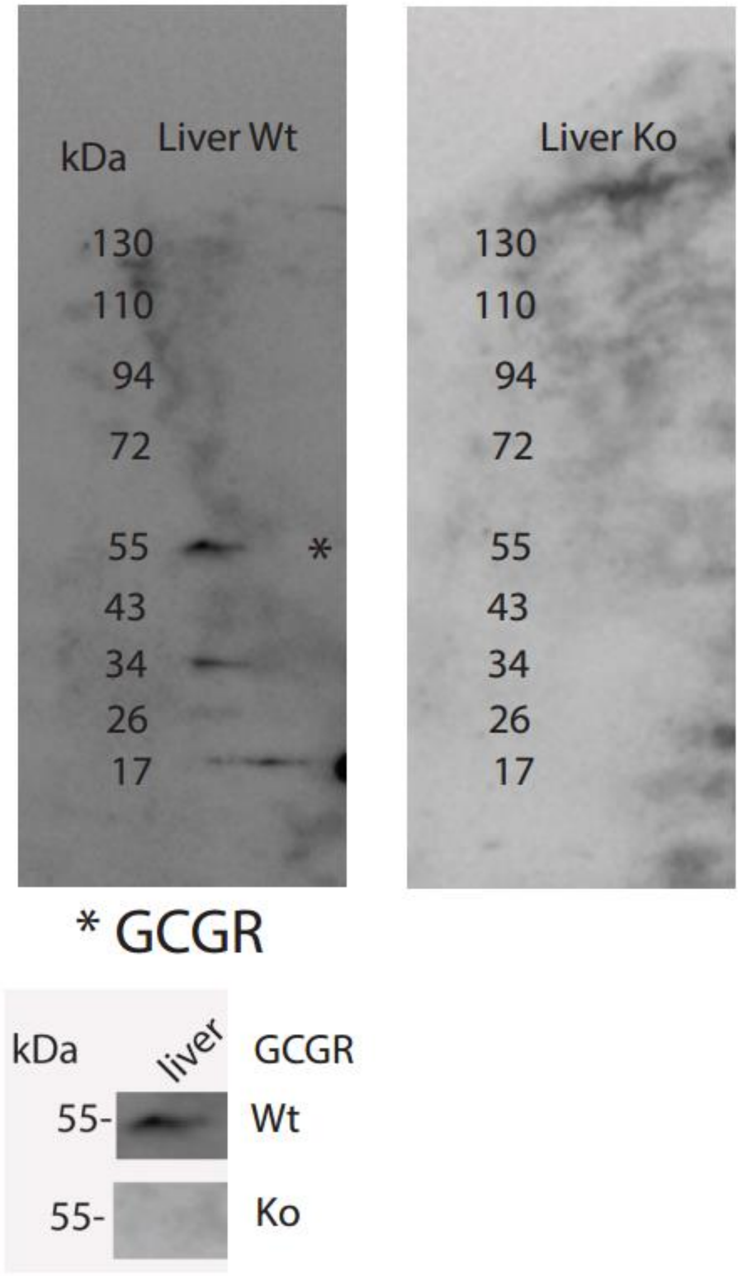


**Supplementary figure 3.** *Uncropped western blotting of lever tissue from Gcgr$^{+/+}$ and Gcgr$^{-/-}$ mice using antibody no. 11*

### Supplementary table 2

**Supplementary table 2**. *Number of samples per tissue used in figure 9. Arranged alphabetically according to tissue name.*

| Tissue name in GTEx | Samples were taken from | Number of GTEx samples |
|---|---|---|
| Adipose Tissue | Subcutaneous & visceral | 406 |
| Adrenal Gland | Adrenal Gland | 56 |
| Blood Vessel | Aorta (Artery), Coronary (Artery), Tibial (Artery) | 409 |
| Brain | Amygdala, Anterior cingulate cortex (BA24), Caudate, Cerebellar Hemisphere, Cerebellum, Cortex, Frontal Cortex (BA9), Hippocampus, Hypothalamus, Nucleus Accumbens, Putamen, Spinal cord (cervical c-1), & Substantia nigra | 1845 |
| Colon | Sigmoid & Transverse | 163 |
| Esophagus | Gastroesophageal Junction, Mucosa, Muscularis | 350 |
| Heart | Atrial Appendage & Left Ventricle | 321 |
| Kidney | Cortex, Medulla | 48 |
| Liver | Liver | 121 |
| Lung | Lung | 213 |
| Muscle | Skeletal | 280 |
| Nerve | Tibial | 217 |
| Pancreas | Pancreas | 38 |
| Pituitary | Pituitary | 207 |
| Prostate | Prostate | 80 |
| Salivary Gland | Minor Salivary Gland | 46 |
| Skin | Not Sun Exposed (Suprapubic) & Sun Exposed (Lower leg) | 639 |
| Small Intestine | Terminal Ileum | 10 |
| Spleen | Spleen | 13 |
| Stomach | Stomach | 43 |
| Testis | Testis | 144 |
| Thyroid | Thyroid | 228 |

### Supplementary table 3

**Supplementary table 3.** *Total number of cells in each cluster used in figure 10. Arranged according to number of cells*

| Liver tissues | | Kidney tissues | |
|---|---|---|---|
| **Cell type (cluster)** | **Number of cells** | **Cell type (cluster)** | **Number of cells** |
| Hepatocytes 1 (1) | 1006 | Tubule cells (1) | 9620 |
| a/b T-cells (2) | 961 | Tubule cells (2) | 7329 |
| Hepatocytes 2 (3) | 909 | Tubule cells (3) | 1974 |
| Inflammatory Macs (4) | 813 | Proximal straight tubule cells (4) | 1267 |
| Hepatocytes 3 (5) | 629 | NK-T cells (5) | 920 |
| Hepatocytes 4 (6) | 603 | Glomerular parital epithelial cells (6) | 734 |
| Plasma cells (7) | 511 | Monocytes (7) | 722 |
| NK-like cells (8) | 488 | Distal tubule cells (8) | 420 |
| y/d T-cells 1 (9) | 464 | Collecting duct principle cells (9) | 161 |
| Non-inflammatory Macs (10) | 379 | B cells (10) | 144 |
| Periportal LSECs (11) | 327 | Collecting intercalated cells (11) | 77 |
| Central Venous LSECs (12) | 306 | | |
| Portal endothelial cells (13) | 211 | | |
| Hepatocytes 5 (14) | 202 | | |
| Hepatocytes 6 (15) | 152 | | |
| Mature B-cells (16) | 129 | | |
| Cholangiocytes (17) | 119 | | |
| y/d T-cells 2 (18) | 105 | | |
| Erythoid cells (19) | 93 | | |
| Hepatic stellate cells (20) | 37 | | |
